# Control theory analysis of dynamic metabolic response elucidates mitochondrial-cytoplasmic coupling and nutrient partitioning

**DOI:** 10.64898/2026.06.28.735091

**Authors:** Xingbo Yang, Daniel J Needleman

## Abstract

Cells adjust their internal circuits in response to changes in their environment. Hence, exposing cells to changing conditions provides a way to probe the intrinsic dynamics of cellular internal circuits. Metabolic networks are examples of such circuits since metabolic fluxes dynamically adjust when environmental conditions are transiently altered. Most existing theoretical frameworks focus on cellular metabolic steady states and do not consider the dynamics of changes in metabolic fluxes. In this work, we applied transfer function analysis from control theory to analyze the changes of NADH oxidative fluxes in the mitochondria and cytoplasm in mouse oocytes in response to dynamical perturbations of oxygen depletion and recovery. We observed an overshoot of NADH oxidative flux in the cytoplasm upon oxygen recovery which is absent in the mitochondrial NADH oxidative flux. Metabolic perturbation experiments and transfer function analysis indicate that this cytoplasmic NADH overshoot results from the coupling of the mitochondrial and cytoplasmic NADH cycles. The degree of overshoot is determined by competing timescales associated with the exchange rates of lactate and pyruvate with the media and their interconversion rates catalyzed by lactate dehydrogenase. Applying control theory to the data enables the inference of the exchange and conversion rates of pyruvate and lactate, allowing predictions of the contribution of lactate to mitochondrial respiration. Our work indicates that the oocytes maintain a homeostatic respiration rate across nutrient conditions by modulating the contribution of lactate to mitochondrial respiration.

## Introduction

Cells face challenges from the external environment, which act as inputs to their internal circuits^1^. How well cells respond and adapt to these inputs depends not only on the regulation mechanisms but also the intrinsic timescales of these cellular circuits that determine how fast and robustly cells respond to dynamical changes of the environmental input^2^. Measuring these intrinsic timescales is a critical step towards understanding the dynamics of the cellular circuits and how these timescales are regulated to generate robust response to environmental fluctuations. Metabolic networks are examples of such cellular circuits. The intrinsic timescales of metabolic pathways are determined by kinetic rates, such as nutrient uptake and conversion rates. Kinetic rates relevant to metabolic pathways are difficult to measure, but metabolic fluxes, which usually depend on a combination of these kinetic rates and metabolite concentrations, can be measured using a variety of different approaches^3^. A wide range of studies have investigated the steady-state behaviors of metabolic fluxes. Many experimental techniques, including extracellular flux measurement^4^ and metabolic flux analysis^5^, are well-suited for measuring steady-state metabolic fluxes. Theoretical techniques, such as metabolic control analysis, have provided frameworks to quantify the metabolic control coefficients associated with different enzymatic steps of a metabolic pathway^6^. However, approaches which focus solely on steady-state fluxes provide only limited information on the underlying kinetic rates which ultimately dictate the behaviors of metabolic networks.

Exposing cells to dynamical perturbations with timescales faster or comparable to the intrinsic timescales of the metabolic networks can induce transient responses whose dynamics are dictated by the kinetic rates of the metabolic networks. Thus, studying the transient response of metabolic networks to dynamic perturbations provides a means to investigate the kinetic rates of metabolic networks. Understanding the transient response of systems to dynamic perturbations is the raison d’être of control theory, which is widely applied in engineering^7^. Control theory provides a quantitative framework to study the relation between the input and output of a dynamical system, with the aim of designing controlling modules to robustly perform a well-defined task. This approach contrasts with the initial condition problems typically faced in physics, where it is of interest to study the time-trajectory of the system given an initial condition without imposing any sense of tasks^8^. In biology, however, one can readily define tasks cells must achieve to survive and proliferate. Control theory has therefore been used in systems biology and synthetic biology^9–11^. However, most applications of control theory to cell biological systems have been purely theoretical and the theory’s applicability and predictive power are rarely tested with experimental data. In this work, we apply transfer function analysis from control theory to experimental data in mouse oocytes to elucidate the coupling mechanisms between mitochondrial and cytoplasmic metabolic cycles and nutrient partitioning.

Glucose, pyruvate and lactate are key nutrients which are catabolized to power cellular functions^12^. This energy conversion is achieved through metabolic cycles, including NADH reduction-oxidation (redox) cycles. These metabolic cycles are compartmentalized in the cell, with NADH redox cycles occurring in both the mitochondria and the cytoplasm. Mitochondria are the sites of cellular respiration, in which the breakdown of pyruvate in the tricarboxylate acid cycle (TCA) results in the reduction of mitochondrial NAD+ to NADH. Mitochondrial NADH is then oxidized back to NAD+ through the electron transport chain (ETC), a process which is coupled to pumping protons through the mitochondrial inner membrane. The resulting electrochemical potential drives the synthesis of ATP. The cytoplasm is the site of glycolysis, in which the breakdown of glucose results in the reduction of cytoplasmic NAD+ to NADH. The cytoplasmic NADH can subsequently either be shuttled into mitochondria to support the ETC or oxidized in the cytoplasm back to NAD+ by lactate dehydrogenase, a process which is coupled to the conversion of pyruvate into lactate. NADH is therefore an important intermediate in energy metabolism that participate in both mitochondrial and cytoplasmic metabolic cycles and couples mitochondrial and cytoplasmic metabolism.

While the metabolic pathways involving NADH are well established, their dynamics in response to metabolic perturbations, and physiological and pathological changes of the cells remain poorly understood. In particular, the dynamics of the coupling between mitochondrial and cytoplasmic NADH cycles remain poorly characterized. Metabolic flux analysis from isotope tracing using mass spectrometry has revealed that the saturation of mitochondrial NADH shuttle drives aerobic glycolysis in cancer cells^13^. Saturation of mitochondrial electron transport chain (ETC) by NADH has been shown to contribute to aerobic glycolysis in yeast^14^. Measurement of NADH autofluorescence dynamics has revealed new mechanisms of metabolic regulations. For example, it has been shown that neuronal stimulation induces a transient increase of NADH levels in the mitochondria as a result of the upregulation of TCA cycles by calcium that upregulates mitochondrial metabolism^15^. Autonomous oscillations of NAD(P)H autofluorescence have been revealed in yeast and are shown to coordinate with cell cycle^16^. In recent years, the development of genetically-encoded NADH/NAD+ biosensors^17^ and NADH oxidases^18^ have enabled the study of the dynamics of NADH redox cycles in mitochondria and cytoplasm. It has been shown that enhancing NADH oxidation in the cytoplasm ameliorates proliferative and metabolic defects caused by an impaired mitochondrial ETC in tissue culture cells^18^, highlighting the importance of the coupling between mitochondrial and cytoplasmic NADH cycles. These results highlight the rich dynamics of NADH redox cycles in the cell in a compartment-dependent manner. Despite the emergence of technology to measure NADH levels with optical resolution, there is a lack of techniques to measure metabolic fluxes through NADH redox cycles in different metabolic compartments in living cells. This significantly limits our understanding of how the dynamics of metabolic pathways are regulated in space and time.

We have previously developed a technique to measure the NADH oxidation rate in mitochondria, which is proportional to the mitochondrial ETC flux, by combining fluorescence lifetime imaging (FLIM) of NADH with biophysical modeling of NADH redox cycles^19^. This technique enables the measurement of ETC fluxes with optical resolution in living cells in a label-free manner. NADH is autofluorescent with a two-photon excitation wavelength at 750nm and emission peak at 460nm that separates itself from other autofluorescent signals in the cell^20^. This autofluorescence enables label-free imaging of NADH levels in living cells. Since NADH is a cofactor, it exists in the cell in two distinct forms: either free in solution or enzyme-bound. When NADH is bound to enzymes, its fluorescence lifetime, defined as the average time the fluorophore spends in the excited state before decaying back to the ground state, increases^21^. Fluorescence lifetime imaging microscopy (FLIM) is a quantitative imaging method that measures fluorescence lifetimes of fluorophores. FLIM of NADH can not only be used to measure the lifetimes of free and bound NADH, but also their respective abundance. It has been shown through extensive research that lifetimes of NADH and the relative abundance of free and bound NADH are correlated with metabolic states of the cell through embryo development^22,23^, cancer progression^24^ and cell differentiation^25^. We have developed a NADH redox model which quantitatively relates the free and bound NADH levels to mitochondrial ETC flux. Using this technique, we have revealed a remarkable phenomenon of flux homeostasis in mouse oocytes, where perturbations of nutrient supply and energy demand of the oocytes do not change mitochondrial ETC flux despite a significant impact on mitochondrial NADH levels^19^. This result calls for a quantitative understanding of how metabolic fluxes are controlled in cells.

Here, we use mouse oocytes arrested at meiosis II (MII) as a model system to study flux control of NADH redox cycles. Meiosis II arrested mouse oocytes are in steady state with a constant metabolic activity, providing an ideal system to investigate perturbations of metabolic activities from a steady baseline. Unlike differentiated somatic cells, mouse oocytes do not perform glycolysis and thus rely exclusively on mitochondrial respiration to produce energy^26^. During oocyte development, oocyte mitochondrial respiration is supported by pyruvate, which is produced in the surrounding cumulus cells and transported through gap junctions into the oocyte^27^. Without pyruvate, mouse embryos cannot develop beyond the two-cell stage^28^. Lactate has been traditionally considered as a metabolic waste from fermentation, but it is increasingly appreciated that it can serve as a major circulating carbohydrate fuel in mammals^29^. A significant level of lactate is present in the oviduct and increasing lactate was shown to decrease pyruvate uptake at the single-cell stage of the mouse embryo^30^, suggesting lactate regulates pyruvate uptake, but the mechanism is unknown. Recent work has revealed that oocytes also consume internal stores of fatty acids^31^. Since a variety of nutrients are involved in oocytes metabolism, we asked how oocytes partition these different nutrients to maintain a homeostatic mitochondrial respiration rate in response to changes in external nutrient conditions.

In this work, we used the coarse-grained NADH redox model to infer the NADH oxidative fluxes from FLIM measurements of NADH in both the mitochondrial and cytoplasmic compartments of mouse oocytes. We observed an overshoot in the cytoplasmic NADH oxidative flux in response to acute oxygen recovery after oxygen depletion. In contrast, the mitochondrial NADH oxidative flux does not display such an overshoot. Inhibition experiments demonstrate that the overshoot dynamics depends on the coupling between mitochondrial and cytoplasmic NADH cycles through the conversion of lactate to pyruvate by lactate dehydrogenase and mitochondrial pyruvate uptake. We use transfer function analysis from control theory to quantitatively explain the flux overshoot based on a competition between the intrinsic timescales of the interconversion between pyruvate and lactate and their exchange with the media. Applying the control theory analysis to the experimental data enables the inference of the contribution of lactate to mitochondrial respiration across a wide range of nutrient conditions. Our work indicates that mouse oocytes maintain a homeostatic respiration rate by modulating the contribution of lactate to mitochondrial respiration in response to changes of external pyruvate and lactate concentrations.

## Results

### Oxygen perturbation induces distinct responses of NADH levels in mitochondria and cytoplasm

We cultured matured mouse oocytes in AKSOM media containing pyruvate, lactate and glucose. We measured oocyte metabolism by imaging NADH using FLIM (Figure 1a). To study the regulation of NADH redox cycles, we sought to determine the dynamics of the response of NADH to a metabolic perturbation. Oxygen is required for mitochondrial respiration and acts as the electron acceptor for NADH. We therefore perturbed NADH dynamics by changing oxygen concentrations in the media. We cultured oocytes in media with 50µM oxygen for 10 mins, and depleted oxygen continuously from 50µM to close to 0µM over 30 mins. To test if the response to this perturbation is reversible, we quickly returned the oxygen level back to 50µM and continued imaging NADH for an additional 20 mins. We observed a dynamical change of NADH intensities in the mouse oocytes in response to oxygen depletion and recovery (Figure 1a). To quantify the NADH intensity changes, we first segmented mitochondria from cytoplasm. Given that NADH concentration is higher in mitochondria than cytoplasm, we segmented mitochondria by identifying brighter NADH pixels using the pixel classification algorithm in ilastik^32^ (Figure 1b). We tested the accuracy of the segmentation by comparing the NADH-based segmentation against MitoTracker-based segmentation. We obtained over 80% accuracy in distinguishing photons coming from mitochondria over cytoplasm^19^. We therefore used only NADH for segmentation without the usage of MitoTracker, rendering our imaging label-free. After the segmentation, we quantified the NADH intensity changes in mitochondria and cytoplasm in response to changes in oxygen levels in the media (Figure 1c). We observed an increase of NADH level in the mitochondria and a decrease of NADH level in the cytoplasm when oxygen drops. Upon oxygen recovery, mitochondrial NADH level drops monotonically back to the initial value, while cytoplasm NADH level overshoots before returning to the initial value (Figure 1c). The full recovery of NADH level suggests that there is no irreversible damage to the oocyte during the oxygen drop. The increase of mitochondrial NADH with decreasing oxygen level is expected due to mitochondrial NADH piling up at lower oxygen levels, since in the absence of oxygen NADH cannot be oxidized to NAD+ and therefore accumulates in the mitochondria. However, it is unclear why cytoplasmic NADH level responds to changes in oxygen levels because NADH redox cycles in the cytoplasm do not depend directly on oxygen. The overshoot in cytoplasmic NADH in response to oxygen recovery is particularly surprising because there is no overshoot in the mitochondrial NADH level. This cytoplasmic NADH overshoot and the subsequent relaxation could encode information about the intrinsic rates of the underlying metabolic pathways coupled to NADH redox cycles in the cytoplasm. We next explore what these metabolic pathways are.

**Figure 1.**
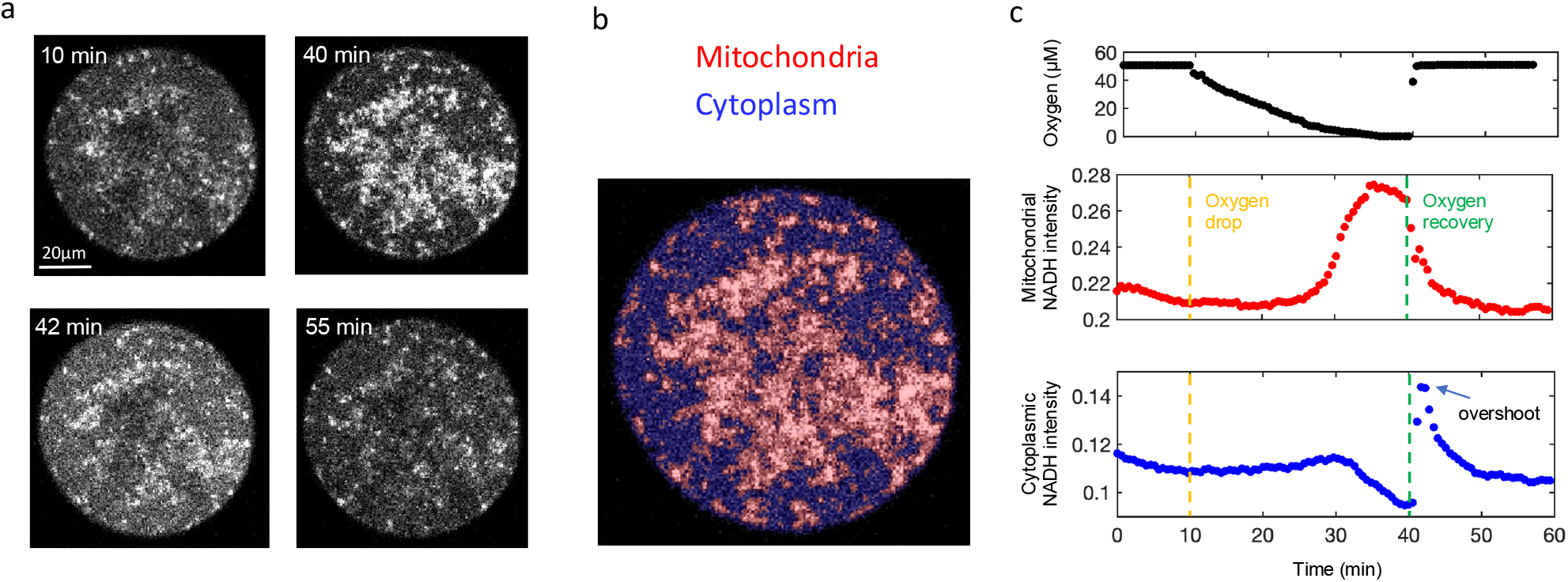
Response of mitochondrial and cytoplasmic NADH intensities to oxygen depletion and recovery. a) Intensity images of NADH in MII mouse oocytes at different time points during an oxygen drop and recovery. b) Segmentation of mitochondria from cytoplasm from pixel classification based on the NADH image. c) Time traces of oxygen level in the media (top), mitochondrial NADH intensity (middle) and cytoplasmic NADH intensity (bottom) during oxygen drop and recovery experiment for a single oocyte.

### Lactate dehydrogenase converts lactate to pyruvate and reduces NAD+ to NADH in the cytoplasm

To better understand the observed dynamic response of NADH levels to modulating oxygen, we next investigated the processes which determine NADH levels in the cytoplasm. In somatic cells, cytoplasmic NADH is produced from NAD+ through glycolysis and oxidized back to NAD+ by lactate dehydrogenase (LDH) or by malate dehydrogenase (MDH)^12^. This forms a complete NADH redox cycle. While LDH is known to be a regulator of redox metabolism in mouse oocytes, it is unclear what completes the cytoplasmic NADH redox cycle in these oocytes^33^ since they exhibit negligible glycolysis^26^. We sought to further clarify the contribution of LDH to cytoplasmic NADH levels in mouse oocytes. We inhibited LDH using 10mM of sodium oxamate and observed a significant decrease of cytoplasmic NADH intensity in the oocytes (Figure 2a). The decrease of NADH level upon LDH inhibition suggests that the net reaction of LDH is to convert NAD+ to NADH. This is in contrast to what has previously been observed in highly glycolytic somatic cells, where oxamate induces an increase of cytoplasmic NADH/NAD^+^ ratio^34^. We therefore concluded that LDH operates in reverse direction in the oocytes compared to glycolytic somatic cells, i.e. LDH converts lactate to pyruvate and reduces NAD+ into NADH in mouse oocytes (Figure 2a). We next asked what pathway oxidizes NADH back to NAD+ to complete the redox cycle in the cytoplasm. Since the malate-aspartate shuttle is known to oxidize cytoplasmic NADH in glycolytic somatic cells, we next investigated the role of the malate-aspartate shuttle in mouse oocytes by inhibiting it using 11mM Aminooxyacetic acid (AOA). This perturbation increases the cytoplasmic NADH level, suggesting that the malate-aspartate shuttle plays a significant role in oxidizing NADH to NAD+ in mouse oocytes (Figure 2b). Taken together, these results indicate that the cytoplasmic NADH redox cycle in mouse oocytes consists of net NAD+ reduction by LDH and NADH oxidation by the malate-aspartate shuttle.

**Figure 2.**
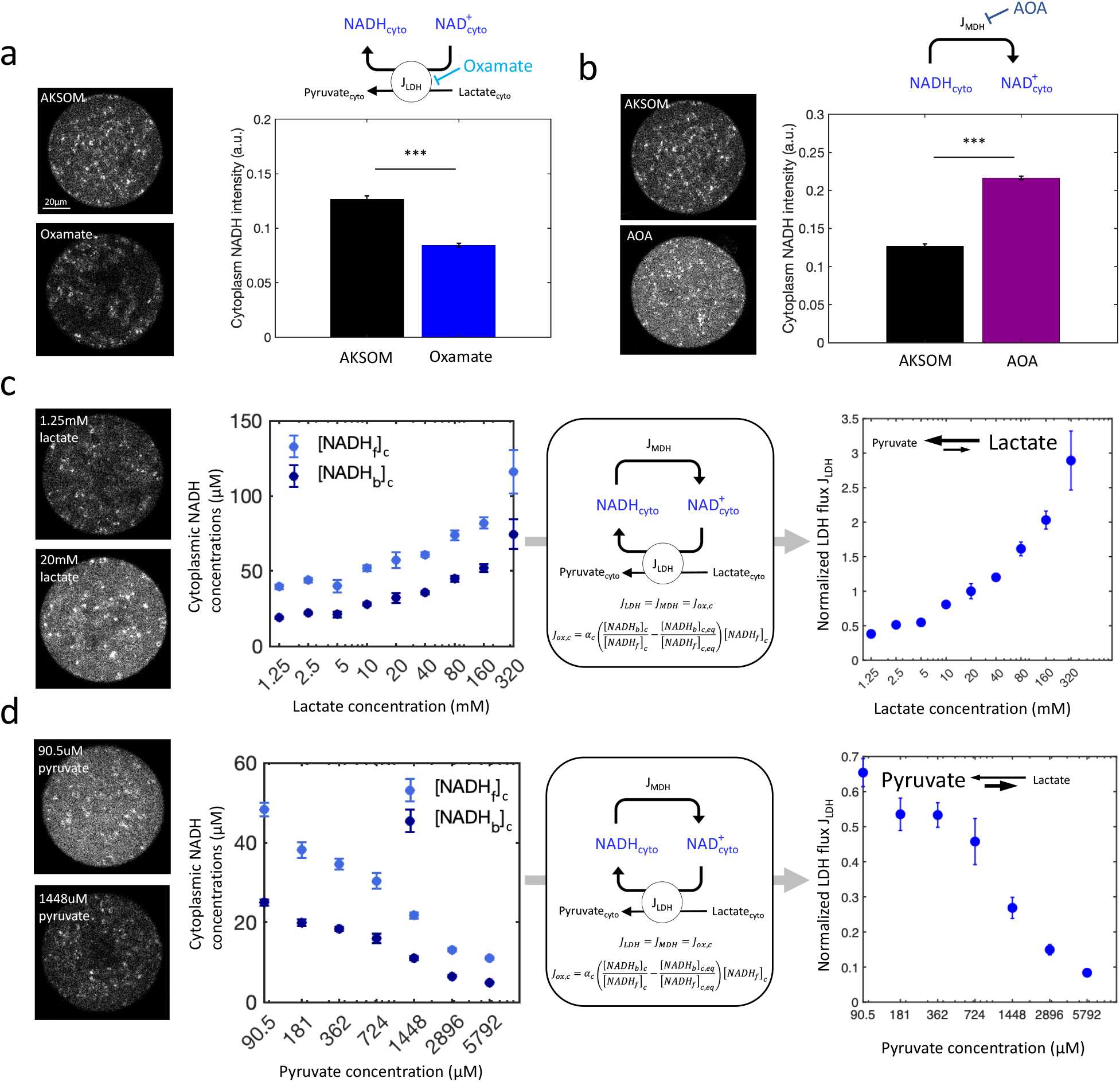
Characterization of cytoplasmic NADH cycles in mouse oocytes. a) Cytoplasmic NADH intensity before and after the addition of 10mM of sodium oxamate that inhibits lactate dehydrogenase (LDH). n=48 for AKSOM and n=24 for oxamate. b) Cytoplasmic NADH intensity before and after the addition of 11mM of AOA that inhibits the malate-aspartate shuttle. n=48 for AKSOM and n=18 for AOA. c) Cytoplasmic free and bound NADH concentrations, [*NADH*]_*c*_ and [*NADH*_*b*_]_*c*_, as a function of lactate concentrations in the media (left). The NADH redox model enables the inference of flux through LDH, *J*_*LDH*_, from the concentrations of cytoplasmic free and bound NADH concentrations (middle). The normalized inferred flux through lactate dehydrogenase as a function of lactate concentrations in the media (right). n=18,18,18,18,30,12,12,12,6 for external lactate concentration from 1.25mM to 320mM, respectively, while fixing external pyruvate concentration at 181 µM. The fluxes are normalized to J_ldh_ at external lactate concentration of 20mM and pyruvate concentration of 181 µM. d) Cytoplasmic free and bound NADH concentrations as a function of pyruvate concentrations in the media (left). The NADH redox model enables the inference of flux through LDH from the cytoplasmic concentrations of free and bound NADH concentrations (middle). The normalized inferred flux through lactate dehydrogenase as a function of pyruvate concentrations in the media (right). n=8,7,8,8,8,12,12 for external pyruvate concentration from 90.5µM to 5792µM, respectively, while fixing external lactate concentration at 10mM. The fluxes are normalized to J_ldh_ at external lactate concentration of 20mM and pyruvate concentration of 181 µM. Error bars denote standard error of the mean across individual oocytes. n denotes number of oocytes.

To further explore the impact of LDH flux in mouse oocytes, we titrated pyruvate and lactate concentrations in the media and used FLIM to measure the corresponding changes of the concentration of NADH bound to enzymes in the cytoplasm, [*NADH*_*b*_]_*c*_, and the concentration of free NADH in the cytoplasm, [*NADH*_*f*_]_*c*_ (Supp Mat Figure 1S and Methods). We observed increases of both free and bound cytoplasmic NADH concentrations with the increase of lactate concentrations (Figure 2c), consistent with the conversion of lactate to pyruvate being accompanied by the reduction of NAD+ to NADH. In contrast, we observed a decrease in the concentrations of free and bound cytoplasmic NADH with increasing pyruvate concentration (Figure 2d), consistent with the conversion of pyruvate to lactate being accompanied by the oxidation of NADH to NAD+. To further test the proposed direction of LDH flux in mouse oocytes (i.e. a net conversion of lactate to pyruvate and of NAD+ to NADH), we used a previously established coarse-grained NADH redox model in conjunction with our FLIM measurements to infer how the flux through LDH varies as a function of lactate and pyruvate concentrations. In short, this model considers a general NADH redox cycle involving an arbitrary number of oxidases and reductases, whose kinetics can obey arbitrary rate laws and may have nonlinear dependencies on metabolite concentrations, enzyme concentrations, and other factors^19^. We coarse-grained all the oxidases into a single effective oxidase and all the reductases into a single effective reductase, while preserving the total NADH oxidative and reductive fluxes through the cycle. The model includes the binding and unbinding of free NADH to the effective oxidase and reductase, the oxidation of NADH to NAD+ by the oxidase and the reduction of NAD+ to NADH by the reductase. The key assumption in our analysis is that the NADH redox cycle is at steady state, where the concentrations of free and enzyme-bound NADH do not change over time. At steady state, the NADH oxidation flux, *J*_*ox*_, is balanced by the net binding and unbinding flux of free NADH to the effective oxidase, which can be calculated as a function of the free and bound concentrations of NADH and the coarse-grained binding and unbinding rates of NADH:

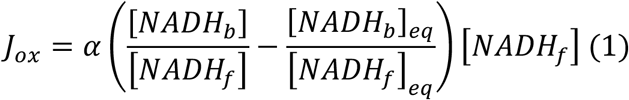

The proportional factor *α* is a function of the unbinding rates of NADH from the oxidase and reductase, while the equilibrium ratio of bound and free NADH concentrations [*NADH*_*b*_]_*eq*_/ [*NADH*]_*eq*_ is a function of both binding and unbinding rates of NADH from the oxidase and reductase^19^. Since the concentrations of free NADH, [*NADH*_*f*_], and bound NADH, [*NADH*_*b*_], can be measured with FLIM, Eq (1) can be used to infer the NADH oxidation flux, *J*_*ox*_. This procedure was originally used to infer NADH oxidation flux in the mitochondria, which is equivalent to inferring the electron transport chain (ETC) flux^19^. However, the coarse-grained NADH redox model is not limited to application in mitochondria because the model construction does not involve any compartment-specific assumptions: Eq (1) can be applied to NADH redox cycles in any cellular compartments where NADH redox cycles exist and operate at steady state, irrespective of the details of the enzymes involved. Since the cytoplasmic NADH redox loop is coupled to the conversion between lactate and pyruvate conversion by LDH, applying Eq (1) to cytoplasmic NADH enables inference of the LDH flux.

Using Eq (1) to infer fluxes requires knowledge of the proportionality factor, *α*, and the equilibrium ratio of bound and free NADH concentrations [*NADH*_*b*_]_*eq*_/[*NADH*_*f*_], which will in general be different in the mitochondria, with *α*_*m*_ and [*NADH*_*b*_]_*m,eq*_/[*NADH*_*f*_], and the cytoplasm, with *α*_*c*_ and [*NADH*_*b*_]_*c,eq*_/[*NADH*_*f*_]. *α*_*m*_ has been verified to be a constant for mitochondria in mouse oocytes, independent of oxygen concentration, and takes a value of *α*_*m*_ = 5.4 ± 0.2*s*^−1 19^ We assume *α*_*c*_ is also a constant throughout the cytoplasm of mouse oocytes. While measuring the precise value of the cytoplasmic NADH flux requires knowledge of the value of *α*_*c*_, Eq (1) can still be used to quantitively infer the relative changes in cytoplasmic NADH flux even if the value of *α*_*c*_ is not known. The equilibrium ratio of bound and free NADH concentrations in mitochondria, [*NADH*_*b*_]_*m,eq*_/[*NADH*_*f*_], can be determined by using FLIM to measure the ratio of the concentration of bound NADH in mitochondria, [*NADH*_*b*_]_*m*_, to that of free NADH in mitochondria, [*NADH*_*f*_]_*m*_, under a condition when there is no oxidative flux of NADH in mitochondria. Since mitochondrial oxidative flux requires the electron transport chain, which in turn requires oxygen, we measured [*NADH*_*b*_]_*m,eq*_/[*NADH*_*f*_]_*m,eq*_ at close to zero oxygen level, giving a value of 0.36 ± 0.01 in AKSOM (Figure 2S d). Similarly, the equilibrium bound ratio in the cytoplasm, [*NADH*_*b*_]_*c,eq*_/[*NADH*_*f*_]_*c,eq*_, can be determined by using FLIM to measure the ratio of the concentration of bound NADH in the cytoplasm, [*NADH*_*b*_]_*c*_, to that of free NADH in the cytoplasm, [*NADH*_*f*_]_*c*_, under a condition when there is no oxidative flux of NADH in the cytoplasm. Suppressing cytoplasm NADH oxidative flux by inhibiting LDH with 10mM of sodium oxamate during oxygen depletion gives an equilibrium bound ration in the cytoplasm, 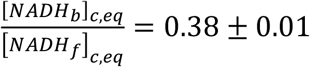, which is taken to be the lowest value of [*NADH*_*b*_]_*c*_ /[*NADH*_*f*_]*_c_* during the oxygen drop in the presence of oxamate in AKSOM (Supp Mat Figure 3S b).

**Figure 3.**
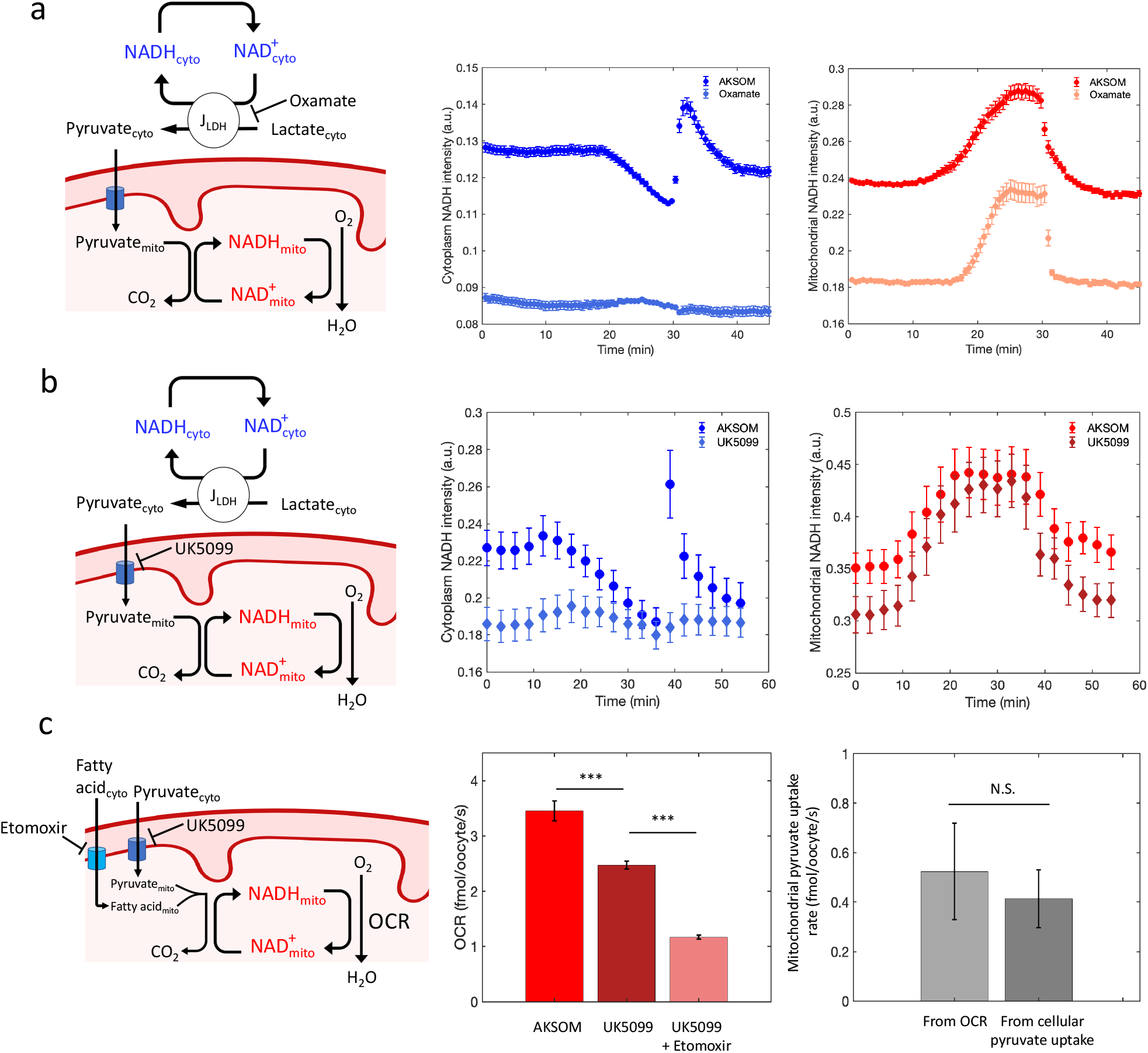
Coupling of cytoplasmic and mitochondrial NADH cycles. a) Schematic showing the inhibition of lactate dehydrogenase using oxamate (left). Response of cytoplasmic NADH intensity to oxygen drop and recovery before and after the addition of 10mM oxamate (middle). Response of mitochondrial NADH intensity to oxygen drop and recovery before (AKSOM) and after (oxamate) the addition of 10mM oxamate (right). n=68 for AKSOM and n=20 for oxamate. b) Schematic showing the inhibition of mitochondrial pyruvate uptake using 5µM of UK5099 (left). Response of cytoplasmic NADH intensity to oxygen drop and recovery before (AKSOM) and after (UK5099) the addition of 5uM of UK5099 (middle). Response of mitochondrial NADH intensity to oxygen drop and recovery before and after the addition of 5µM of UK5099 (right). n=16 for AKSOM and n=16 for UK5099. c) Schematic showing the inhibition of mitochondrial pyruvate uptake using 5µM of UK5099 and fatty acid transport into the mitochondria with 100µM of etomoxir (left). Oxygen consumption rate of the oocyte in control condition (AKSOM), UK5099 and UK5099 plus etomoxir (middle). N=4,2,2 for AKSOM, UK5099 and UK5099 plus etomoxir OCR measurements, respectively. Comparison of mitochondrial pyruvate uptake rate inferred from OCR and from cellular pyruvate uptake in the absence of lactate in the media (right). N=2 for cellular pyruvate uptake measurement. Error bars denote standard error of the mean across individual oocytes for Figure 3a-b, while error bars denote standard error of the mean across batches of experiment for Figure 3c. n denotes number of oocytes. N denotes number of batch of experiments. Two-sample t-test is performed for statistical testing.

At steady state, the NADH oxidation flux in the cytoplasm, *J*_*ox,c*_, is equal to the flux through LDH, *J*_*LDH*_, and, if the cytoplasmic NADH redox cycles is driven by LDH and malate dehydrogenase, this must also equal the flux through malate dehydrogenase, *J*_*MDH*_: i.e. *J*_*ox,c*_ = *J*_*LDH*_ = *J*_*MDH*_. This is true regardless of the direction of the LDH flux because at steady state the NADH oxidation flux balances the NADH reduction flux. Using the concentrations of cytoplasmic free and bound NADH measured by FLIM during the titration of pyruvate and lactate, [*NADH*_*f*_]_*c*_ and [*NADH*_*b*_]_*c*_, (Figure 2 c,d left panel) and Eq (1), we inferred the LDH flux as a function of lactate and pyruvate concentrations (Figure 2 c,d right panel). The inferred LDH flux increases with increasing lactate concentration and decreases with increasing pyruvate concentration, suggesting that lactate is a reaction substrate and pyruvate is a reaction product. Hence, we concluded that the net reaction of LDH is converting lactate to pyruvate and reducing NAD+ to NADH in the oocytes. This is consistent with both i) the observed changes in NADH level upon LDH inhibition (Figure 2a) and ii) that the most abundant isoform of LDH in mouse oocytes is LDH-B, which is the isoform of LDH that most preferably converts lactate to pyruvate^35^.

### Cytoplasmic NADH cycle is coupled to mitochondrial NADH cycle through lactate dehydrogenase and mitochondrial pyruvate uptake

Having revealed the direction of the LDH flux in mouse oocytes, with net conversion of lactate to pyruvate accompanied by the reduction of cytoplasmic NAD+ to NADH, we next investigated the contribution of LDH to the response of cytoplasmic NADH to changes in oxygen levels. We inhibited LDH with 10mM of sodium oxamate and measured cytoplasmic NADH intensity during oxygen drop and recovery (Figure 3a left). The addition of oxamate lowered NADH levels in both the cytoplasm and the mitochondria, but while the overall dynamics of NADH changes in mitochondria remained similar to control (Figure 3a right), the dynamics of NADH changes in the cytoplasm was greatly diminished, with no significant dip or overshoot in response to oxygen drop and recovery (Figure 3a middle).

This result suggests that LDH is required for the cytoplasmic NADH response to oxygen drop and recovery. Since LDH converts lactate to pyruvate and pyruvate is taken up by mitochondria to sustain respiration, we hypothesized that mitochondrial pyruvate uptake couples the cytoplasmic and mitochondrial NADH cycles. To test this hypothesis, we inhibited the mitochondrial pyruvate carrier MPC with 5µM of UK5099 and repeated the oxygen drop experiment (Figure 3b left). The addition of UK5099 diminished the cytoplasmic NADH response to oxygen drop and recovery (Figure 3b middle), while the mitochondrial NADH levels were slightly lowered, their dynamics where largely unaffected by UK5099 (Figure 3b right). The dependency of cytoplasmic NADH’s decrease and overshoot on MPC activity suggests that the cytoplasmic NADH cycle is coupled to the mitochondrial NADH cycle through their mutual use of pyruvate.

The observation that mitochondrial NADH increases in response to decreasing oxygen even after transport of pyruvate into mitochondria is inhibited by UK5099 suggests that pyruvate is not the only source of electrons used to reduce mitochondrial NAD+ to NADH. Fatty acids, which oocytes are known to store and consume, are another source of electrons for reducing NAD+, thereby fueling mitochondrial respiration^31^. We next sought to determine the relative contributions of fatty acids and pyruvate to mitochondrial respiration. We first measured the change of oxygen consumption rate (OCR) after the addition of 5µM UK5099 (Methods), which decreased from 3.45±0.18 fmol/s/oocyte to 2.47±0.07fmol/s/oocyte (Figure 3c middle). Next, we added both UK5099 and an inhibitor of mitochondrial fatty acids transporter CPT-1, etomoxir, to the media (Figure 3c left), resulting in a further decrease in respiration to 1.16±0.03 fmol/s/oocyte (Figure 3c middle). While the simultaneous addition of UK5099 and etomoxir does not completely inhibit the OCR, and the percentage of OCR remaining is similar to that which remains after inhibition of the electron transport chain by rotenone^19^. This suggest that the OCR which remains after addition of UK5099 and etomoxir is non-mitochondrial in origin. Comparing the change of OCR due to inhibiting pyruvate uptake of mitochondria with UK5099 (0.98±0.19 fmol/s/oocyte) and that due to also inhibiting fatty acid uptake of mitochondria using etomoxir (1.31±0.07 fmol/s/oocyte), we concluded that pyruvate and fatty acids each contribute ~50% of mitochondrial respiration.

To further test this estimate of the relative contributions of pyruvate and fatty acids to mitochondrial respiration, we measured the cellular pyruvate uptake rate in the absence of lactate in the media, which was 0.41±0.11 fmol/s/oocyte (Figure 3c right; Methods). To within experimental error, this directly measured cellular pyruvate uptake rate is the same as the mitochondrial pyruvate uptake rate inferred from OCR measurements, which yields 0.52±0.19 fmol/s/oocyte (Figure 3c right), assuming half of the OCR is contributed by pyruvate and a stoichiometry coefficient of oxygen to pyruvate of 3. The agreement between these two measurements supports the conclusion that pyruvate contributes to half of mitochondrial OCR and indicates that pyruvate is not significantly diverted to other pathways.

We have also observed that mitochondrial respiration rate remains unchanged when mitochondrial fatty acid uptake is inhibited by 100µM of etomoxir (Supp Mat Figure 4Sb). This demonstrated that mitochondrial pyruvate uptake can compensate for fatty acid uptake to support mitochondrial respiration. This observation is consistent with the fact that etomoxir alone does not decrease ATP levels in mouse oocytes^31^. On the contrary, inhibiting mitochondrial pyruvate uptake with 5µM of UK5099 decreases OCR, suggesting that fatty acid uptake cannot fully compensate for pyruvate uptake to support mitochondrial respiration (Supp Mat Figure 4Sb).

### Flux through lactate dehydrogenase overshoots after oxygen recovery while mitochondrial pyruvate uptake flux recovers without overshoot

Metabolic activities are characterized by metabolic fluxes. After demonstrating that LDH and mitochondrial pyruvate uptake couple mitochondrial and cytoplasmic NADH cycles, we next investigated how metabolic fluxes through LDH and mitochondrial pyruvate uptake rate respond to oxygen depletion and recovery. Since the carbon fluxes through LDH and mitochondrial pyruvate uptake are coupled to NADH oxidation fluxes, we sought to infer NADH oxidation fluxes in mitochondria and cytoplasm, using the coarse-grained NADH model (i.e. Eq 1), and relate them to carbon fluxes. This entails applying Eq (1) during the time course of oxygen drop and recovery, but Eq (1) is strictly valid at steady state, i.e., when the concentrations of free and enzyme-bound NADH do not change over time. However, if changes in NADH levels occur sufficiently slowly that they can be well approximated as being at steady state, then Eq (1) can still be used even though NADH levels are not constant. This quasi-static regime holds when the rate of change of NADH concentrations is much slower than *J*_*ox*_. During these oxygen drop and recovery experiments, the rate of change of [*NADH*_*f*_]_*m*_ is ~2.5*μM*/*min*, which is much slower than the rapid turnover of NADH in the mitochondrial NADH cycle, 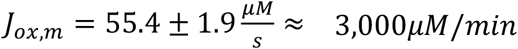^19^. Thus, the mitochondrial NADH dynamics changes quasi-statically during the oxygen drop and recovery. We started by assuming that the cytoplasmic NADH cycle also changes quasi-statically during the oxygen drop and recovery, which could result from slower variation in carbon fluxes (with NADH quasi-statically following the time varying carbon fluxes). We later tested this hypothesis and show that this interpretation is self-consistent (see below).

We first measured the NADH intensity, short and long fluorescence lifetimes and bound fractions in both the cytoplasm and mitochondria from FLIM imaging (Supp Mat Figure 2S). Assuming free and bound NADH molecular brightness is proportional to the short and long fluorescence lifetimes, respectively, and with an *in vitro* calculation of the proportional constant^19^ (Methods), we calculated the absolute concentrations of free NADH and bound NADH in both cytoplasm, [*NADH*_*f*_]_*c*_ and [*NADH*_*b*_]_*c*_, and mitochondria, [*NADH*_*f*_]_*m*_ and [*NADH*_*b*_]_*m*_, in MII mouse oocytes in response to oxygen depletion and recovery (Figure 4b,c left panel). To obtain the NADH oxidation fluxes in mitochondria, *J*_*ox,m*_,and cytoplasm, *J*_*ox,c*_, we applied the NADH redox model that relates NADH oxidation flux to the concentrations of free and bound NADH from Equation 1 to both the mitochondria and cytoplasm (Figure 4b,c, middle panel).

**Figure 4.**
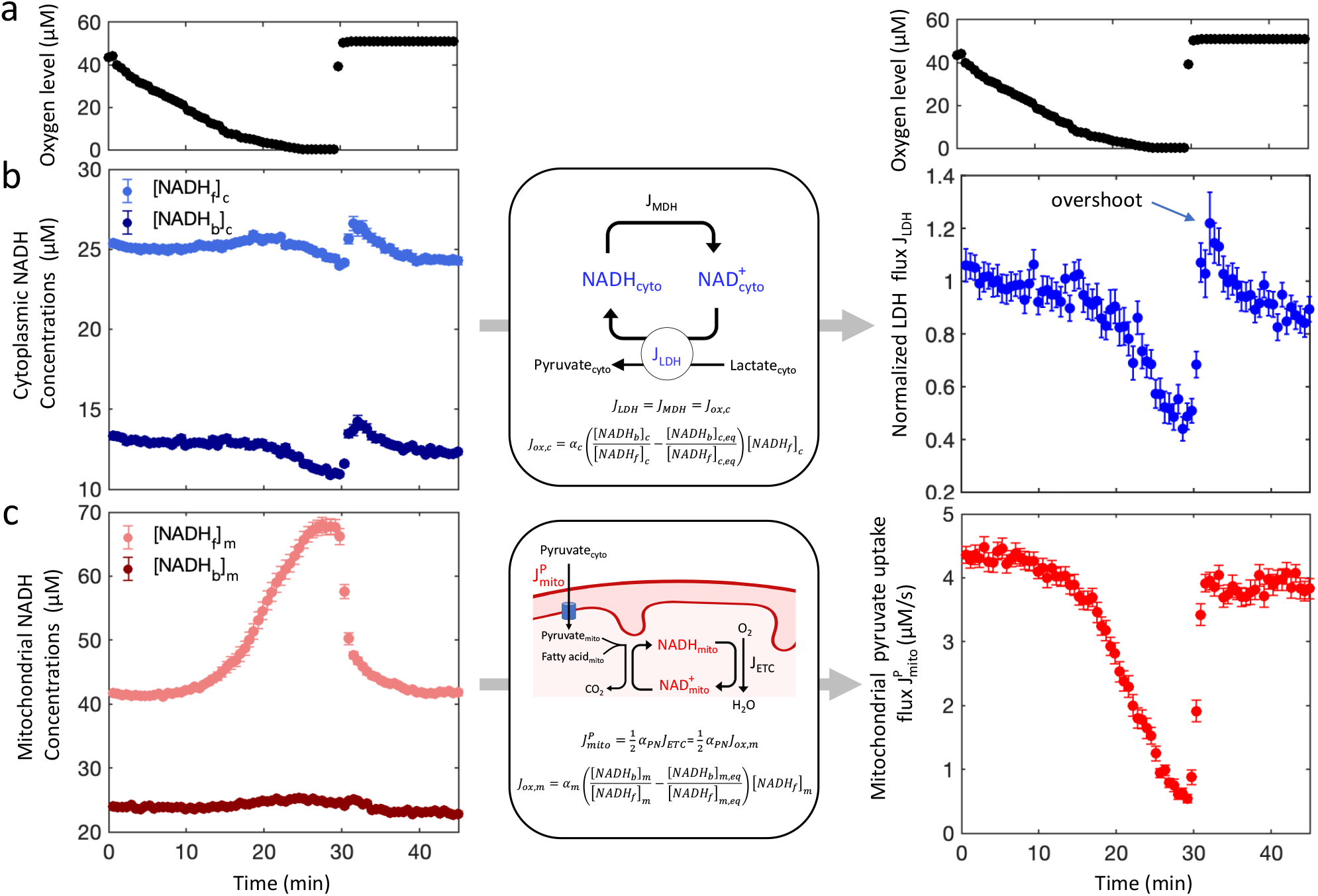
Inferring temporal dynamics of flux through lactate dehydrogenase and mitochondrial pyruvate uptake flux in response to oxygen depletion and recovery. a) Oxygen level in the media during oxygen drop and recovery experiment. b) Cytoplasmic free ([NADH_f_]_c_) and bound ([NADH_b_]_c_) NADH concentrations in response to oxygen drop and recovery (left). NADH redox model that enables the inference of flux through LDH in the cytoplasm (middle). Inferred normalized LDH flux in response to oxygen drop and recovery (right). c) Mitochondrial free ([NADH_f_]_m_) and bound ([NADH_b_]_m_) NADH concentrations in response to oxygen drop and recovery (left). NADH redox model that enables the inference of mitochondrial pyruvate uptake flux (middle). Inferred mitochondrial pyruvate uptake flux in response to oxygen drop and recovery (right). n=68. Error bars denote standard error of the mean across individual oocytes. n denotes number of oocytes.

Next, we related the inferred NADH oxidation fluxes to carbon fluxes. As discussed above, the cytoplasmic NADH oxidative flux equals the flux through LDH: i.e. *J*_*ox,c*_ = *J*_*ldh*_. The inferred oxidative flux of cytoplasmic NADH also equals LDH’s flux of pyruvate because the reduction of one molecule of NAD+ to NADH (which is balanced by oxidation of one NADH back to NAD+) is accompanied by the conversion of one molecule of lactate to pyruvate. Since the proportionality factor, *α*_*c*_, is unknown, it is only possible to use Eq (1) to infer relative changes in LDH flux. The inferred relative flux of cytoplasmic pyruvate from LDH decreases with decreasing oxygen level and, after oxygen recovery, overshoots before relaxing back to a steady-state value (Figure 4b right).

The inhibition of the MPC and fatty acid oxidation, and direct measurements of cellular pyruvate uptake rates, indicate that approximately half of the mitochondrial OCR is accounted for by mitochondrial uptake of pyruvate (Figure 3c), which is equivalent to stating that approximately half of the oxidation of mitochondrial NADH is associated with mitochondrial uptake of pyruvate. If it is assumed that mitochondrial uptake of pyruvate continues to contribute half of the oxidation of mitochondrial NADH throughout the oxygen drop and recovery, then the inferred NADH oxidation flux in the mitochondria in mitochondria, *J*_*ox,m*_, can be converted to an inferred mitochondrial pyruvate uptake flux, 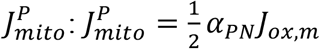 where 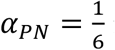 is the stoichiometry factor from detailed knowledge of the TCA and ETC cycles (Methods). The inferred mitochondrial pyruvate uptake rate decreases with decreasing oxygen level and recovers without any overshoot after oxygen recovery (Figure 4c).

### Control theory explains the overshoot of LDH flux from mitochondrial-cytoplasmic coupling and timescales associated with pyruvate and lactate kinetics

The inferred reduction in mitochondrial pyruvate uptake rate with decreasing oxygen level is expected since oxygen is the terminal electron acceptor required to run the electron transport chain. It is less clear why LDH flux responds to oxygen depletion because oxygen is not a direct substrate for the reaction catalyzed by LDH. The observation that the cytoplasmic NADH response to oxygen depletion is diminished after the inhibition of mitochondrial pyruvate transport (Figure 3b middle) indicates that the cytoplasmic NADH response to oxygen is coupled to mitochondria via mitochondrial pyruvate transport. However, it is not obvious how mitochondrial pyruvate transport could lead to an overshoot in LDH flux after oxygen recovery, particularly because such an overshoot is not present in mitochondrial pyruvate uptake rate (compare Figures 4b and 4c).

To gain further insight into the origin of the overshoot, we next developed a kinetic model of pyruvate and lactate metabolism in mouse oocytes. Our minimal model of the dynamics of the concentrations of cytoplasmic pyruvate, *P*_*cyto*_, and lactate, *L*_*cyto*_, contain fluxes from exchange of pyruvate and lactate with the external media, 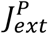 and 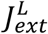, and the interconversion flux via LDH, *J*_*ldh*_, and the consumption of pyruvate by mitochondria, 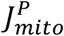, (Figure 5a):

**Figure 5.**
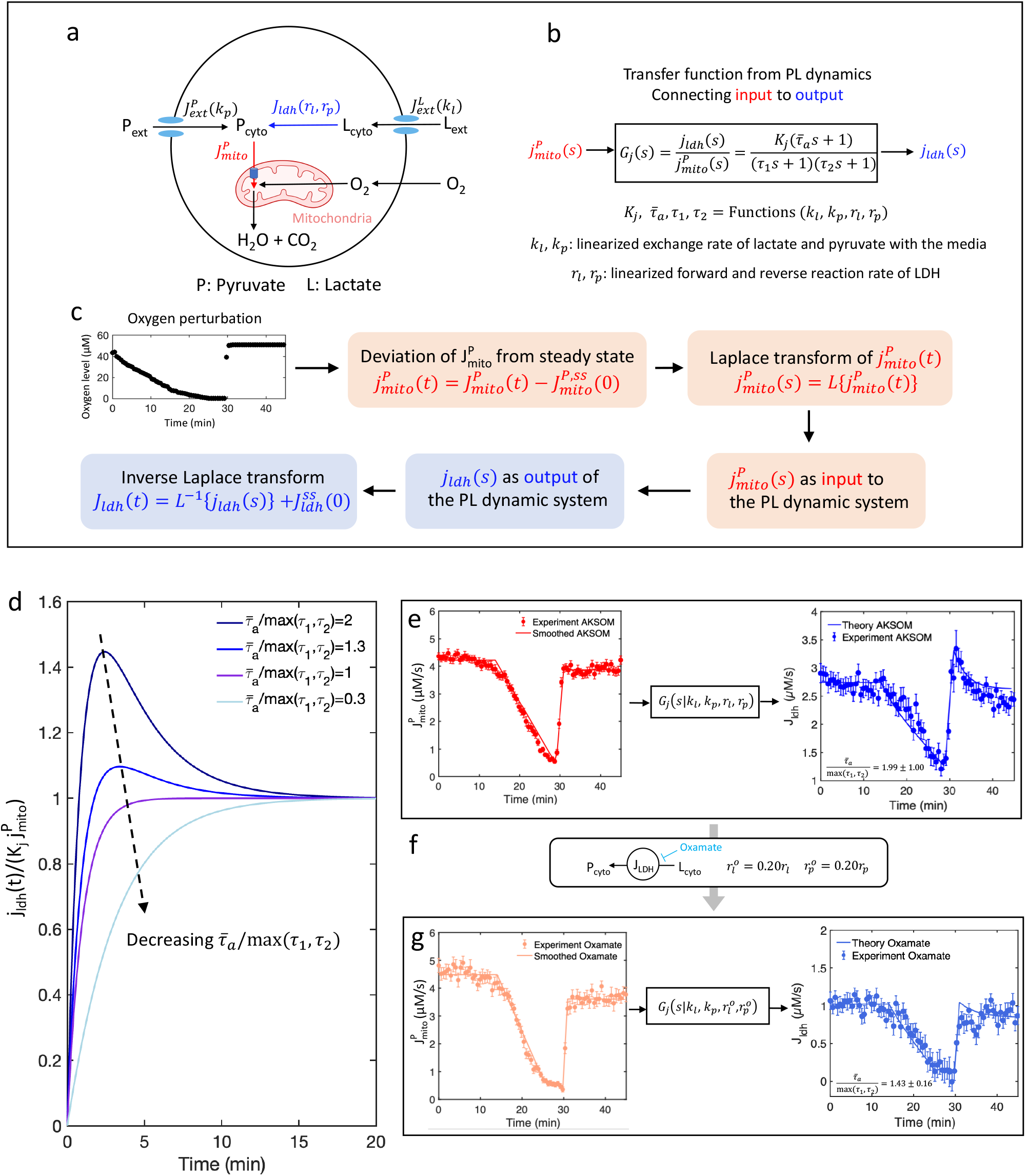
Overshoot in LDH flux is controlled by timescales associated with interconversion rates of pyruvate and lactate and their exchange rates with the media. a) Schematic of the kinetic pyruvate lactate metabolism model. b) Transfer function (G_j_) connecting mitochondrial pyruvate uptake flux (j^P^_mito_) as the input to the flux through lactate dehydrogenase (j_ldh_) as the output. c) Workflow of predicting the response of flux through lactate dehydrogenase to time-varying mitochondrial pyruvate uptake flux using control theory. d) Predicted response of LDH flux in response to a step increase of mitochondrial pyruvate uptake flux at different ratios of the characteristic timescales. e) Numerical solution of the kinetic model for LDH flux in response of oxygen drop and recovery taking smoothed mitochondrial pyruvate uptake flux as the input. f) Reducing pyruvate and lactate conversion rate upon inhibition of LDH with 10mM of sodium oxamate. g) Numerical solution of the kinetic model for LDH flux in response of oxygen drop and recovery after oxamate perturbation taking smoothed mitochondrial pyruvate uptake flux as the input. n=68 for AKSOM and n=20 for oxamate. Error bars denote standard error of the mean across individual oocytes. n denotes number of oocytes.

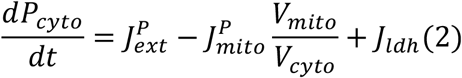

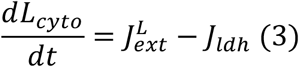

In general, these fluxes could be functions of *P*_*cyto*_, *L*_*cyto*_ and other variables. The fluxes could be either positive or negative depending on the direction of the fluxes. To proceed, we assumed that the oxygen drop and recovery experiment perturbs the cytoplasmic pyruvate and lactate concentrations around their initial steady-state values, such that 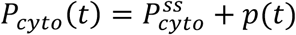 and 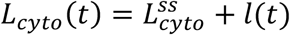. We further assumed that *J*_*ldh*_ is a function of cytoplasmic pyruvate and lactate concentrations while 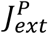 and 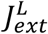 are functions of both cytoplasmic and external pyruvate and lactate concentrations. During the oxygen perturbation, the external pyruvate and lactate concentrations are kept fixed. Hence, we can linearly expand *J*_*ldh*_, 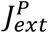 and 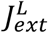 as a function of *p*(*t*) and 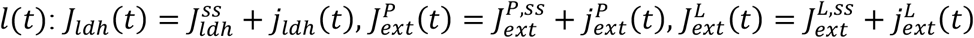, where 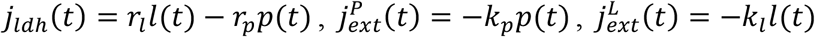. Here, *r*_*l*_, *r*_*p*_, *k*_*l*_ and *k*_*p*_ are related to the first order derivatives of *J*_*ldh*_(*t*), 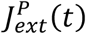 and 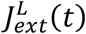 with respect to *P*_*cyto*_(*t*) and *L*_*cyto*_(*t*) evaluated at the steady values 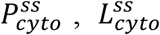 and external pyruvate and lactate concentrations (Supp Mat Eq S8-S9). They are linearized first order rates that characterize the sensitivities of the fluxes to perturbations of the cytoplasmic pyruvate and lactate concentrations around their steady-state values. We also expand the pyruvate uptake flux around its initial steady value 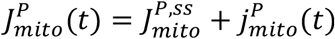. By definition, 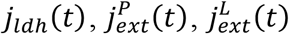, and 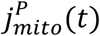 are all zero at *t* = 0, defined as the beginning of the experiment before oxygen drop. Finally, 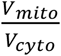 is the ratio between mitochondrial volume and the cytoplasm volume, which is approximately 1 from the NADH fluorescence measurement^19^. Taken together, these steps result in dynamical equations for *p*(*t*) and *l*(*t*) (Supp Mat Eq S6-S7):

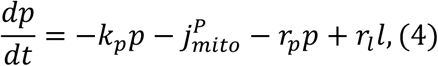

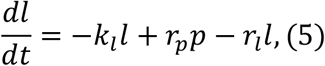

We next sought to use this model to investigate the origin of the overshoot in LDH flux that occurs after oxygen recovery (Figure 4b). We hypothesized that changes in oxygen levels impact mitochondrial respiration, which induces changes in the uptake of pyruvate to mitochondria, 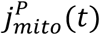. This change in 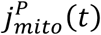 induces changes in cytoplasmic pyruvate, *p*(*t*), and lactate, *l*(*t*), concentrations, which leads to the alterations in the LDH flux, *j*_*ldh*_(*t*). We used the framework of control theory to aid in evaluating the plausibility of this scenario.

A key concept in control theory is the transfer function, which relates a system’s inputs to its outputs^7^. We consider the uptake of pyruvate to mitochondria as the input and the flux of LDH as the output (Figure 5b). More precisely, the transfer function, *G*_j_(*s*), allows the time-dependent deviations in the flux of LDH, *j*_*ldh*_(*s*) to be calculated from the time-dependent deviations in the uptake of pyruvate to mitochondria, 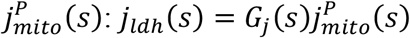, where these functions are in Laplace space. The transfer function in this model can be analytically calculated (Supp Materials Eq S20):

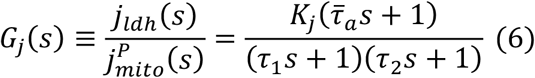

Where 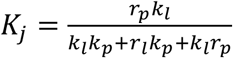 is an overall scale factor that sets the magnitude of the transfer function and *τ*_1_, *τ*_2_, and 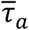 are the three intrinsic timescales that characterize the dynamics of the transfer function, with:

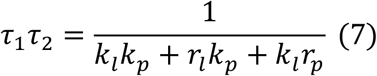

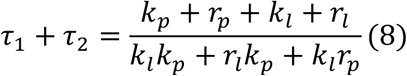

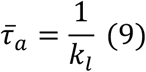

This transfer function can be used to calculate the response of LDH flux dynamics in real time space, *j*_*ldh*_(*t*), for any given dynamics of the mitochondrial uptake flux, *j*^*P*^ (*t*), through inverse Laplace transform (Figure 5c).

We next used this transfer function to investigate if an overshoot in *j*_*ldh*_(*t*) could be produced from oxygen recovery rapidly changing 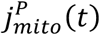, even though 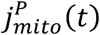 does not exhibit an overshoot. We modeled 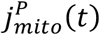 as undergoing an instant change during oxygen recovery going from zero at low oxygen to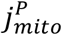 at high oxygen, i.e. 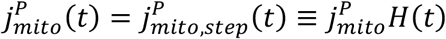, where *H*(*t*) is the Heaviside step function. We use the transfer function to analytically predict the resulting time course of *j*_*ldh*_(*t*) = *j*_*ldh,step*_(*t*), where

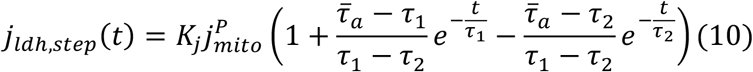

This function exhibits an overshoots if 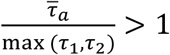, with the magnitude of the overshoot increasing as 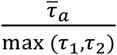 increases (Figure 5d). In this model, both lactate and pyruvate concentrations change monotonically in response to a step change in 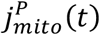, without exhibiting any overshoot themselves, but they do so at different rates (Figure 5Sa). The net fluxes through LDH is due to the difference between the forward flux (i.e. *r*_*l*_*l*(*t*)), which scales with lactate concentration, and the reverse flux (i.e. *r*_*p*_*p*(*t*)), which scales with pyruvate concentration, and the different relaxation rates of pyruvate and lactate can therefore result in an overshoot in *j*_*ldh*_(*t*). Thus, this kinetic model, which only incorporates simple representations of the exchange of lactate and pyruvate with the media, their interconversion by LDH, and the uptake of pyruvate by mitochondria, is sufficient to explain the overshoot in LDH flux due to a sudden change in the uptake of pyruvate by mitochondria.

We next sought to perform a more quantitative comparison between the predictions of the kinetic model and experiments. We solved eqn (4) and (5) numerically with an initial condition of *p* = 0 and *l* = 0, and we used the experimentally inferred deviation of uptake of pyruvate by mitochondria, 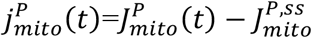 as an input where 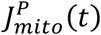 and 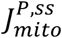 are obtained from Figure 5e, left. For numerical stability, we smoothed 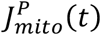 by fitting it with piecewise linear functions (Figure 5e, left). The numerical solution yields the deviation of LDH flux from the initial steady value, *j*_*ldh*_(*t*). We obtained the full LDH flux from 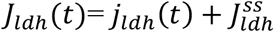. We adjusted the kinetic model’s parameters (*k*_*l*_, *k*_*p*_, *r*_*l*_, *r*_*p*_) and the initial steady state LDH flux 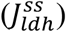 to best fit the full dynamics of experimentally inferred LDH flux, *J*_*ldh*_(*t*) (Figure 5e, right). The kinetic model well fitted the data with *k*_*l*_ = 0.0028 ± 0.0006*s*^−1^, *k*_*p*_ = 0.3 ± 0.1*s*^−1^, *r*_*l*_ = 0.0086±0.004*s*^−1^ and *r*_*p*_ = 0.69 ± 0.06*s*^−1^, providing absolute measures of these parameters. These fitted values give 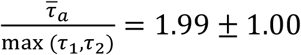, which is greater than 1, as expected since an overshoot in, *j*_*ldh*_(*t*) is observed. We note that these kinetic parameters are only expected to hold for oocytes cultured in AKSOM, since the parameters implicitly depend on the external pyruvate and lactate concentrations and the steady state cytoplasmic pyruvate and lactate concentrations around which the perturbations are performed. Remarkably, the fitting procedure directly gives the best fitting steady-state LDH flux 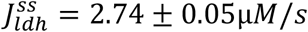 (Figure 5e right). Knowing the absolute 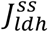 enables us to calibrate the kinetic constant in the NADH redox model for the cytoplasm *α*_*c*_ = 0.81 ± 0.01*s*^−1^, which can be used to infer LDH flux in other conditions.

To further investigate the validity of the kinetic model, we next used it to study how the response of the oocytes was modified by LDH inhibition. We used FLIM and the coarse-grained NADH model to infer how mitochondrial pyruvate uptake rate, 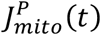, and LDH flux, *J*_*ldh*_(*t*), varied in response to oxygen drop and recovery in the presence of 10mM of sodium oxamate. The presence of oxamate had minimal impact on the inferred 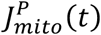 during oxgen drop and recovery, but the inferred *J*_*ldh*_(*t*) was strongly reduced and exhibited a greatly diminished overshoot (Figure 5g and Supp Mat Figure 3S). We again numerically solved Eqn (4) and (5) and used the smoothed experimentally inferred deviation of uptake of pyruvate by mitochondria, 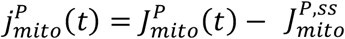 in the presence of oxamate, as an input (Figure 5g, left) and performed a best fit to the experimentally inferred LDH flux, *J*_*ldh*_(*t*) in the presence of oxamate (Figure 5g, right), keeping all kinetic parameters fixed except allowing *r*_*l*_ and *r*_*p*_ that characterize LDH kinetics to change by a multiplicative factor 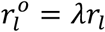 and 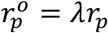. Since we obtained *α_c_* from the AKSOM experiment, the initial steady-state LDH flux in the presence of oxamate is inferred as 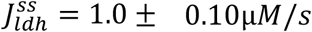, eliminating the need to fit 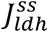. We hence only need to fit the multiplicative factor *λ* to best fit the observed *J*_*ldh*_(*t*) in the presence of oxamate, yielding *λ* = 0.20 ± 0.01, reducing the LDH associated rate constants such that 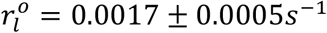 and 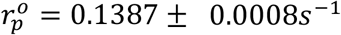, which resulted in an excellent agreement between inferred and predicted LDH fluxes in the presence of oxamate (Figure 5g, right). This change in LDH rate constants leads to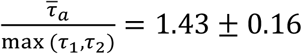 0.16, a reduction which is consistent with the observed decrease in overshoot. Thus, the kinetic model is sufficient to explain the impact of LDH inhibition on the response of oocytes, further supporting its utility.

To understand how the overshoot of LDH flux depends on the kinetic rates of the model, we consider the limit where *r*_*p*_, *k*_*p*_ ≫ *r*_*l*_, *k*_*l*_, which is consistent with the values of the fitted parameters *k*_*l*_ = 0.0028 ± 0.0006*s*^−1^, *k*_*p*_ = 0.3 ± 0.1*s*^−1^, *r*_*l*_ = 0.0086±0.004*s*^−1^ and *r*_*p*_ = 0.69 ± 0.06*s*^−1^. In this limit, the overshoot occurs when 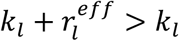, where 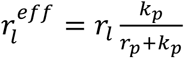 (See Supp Mat eqns S25-S28). Therefore, the overshoot is controlled by the conversion rate of lactate to pyruvate scaled by the fraction of pyruvate that exchanges with the media.

Intuitively, the transient overshoot of cytoplasmic NADH is induced by non-steady state dynamics of pyruvate and lactate concentrations as predicted by the kinetic model. Specifically, the rate of change of intracellular pyruvate concentrations in response to the recovery of oxygen level is different from the rate of change of the lactate concentrations because pyruvate is directly taken up by the mitochondria. This asymmetry of response rates leads to an enhanced difference between the forward and reverse reaction rates of LDH, resulting in an overshoot of the net flux (Supp Mat Figure 5S).

For generality, we also consider the case when cytoplasmic NADH redox cycle is not at steady state. In this case, we developed a generalized kinetic model that directly predicts the overshoot of NADH concentrations (Supp Mat Eqns S33-35) to the change of mitochondrial pyruvate uptake flux without the need to infer LDH flux (Eqn.S40). This model involves an additional timescale of the NADH oxidation and reduction. The generalized kinetic model reduces to the kinetic model in the limit when the NADH oxidation and reduction dynamics is faster than the pyruvate and lactate dynamics.

We observed that the absolute value of *J*_*ldh*_(*t*) decreases after oxamate perturbation, but the mitochondrial pyruvate uptake flux 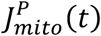 remains unchanged (Figure 5e and g left). This observation is consistent with the phenomenon of flux homeostasis that we discovered in our earlier work^19^ and subsequently also observed in growing yeast^14^, where the mitochondrial oxygen consumption rate is insensitive to nutrient perturbations despite significant changes in NADH concentrations.

### Oocytes maintain a homeostatic mitochondrial respiration rate by modulating the contribution of lactate to respiration

The homeostasis of mitochondrial respiration rate after LDH inhibition suggests redistribution of nutrient fluxes in response to perturbations of nutrient supplies. Pyruvate and lactate are both important nutrients for oocytes and early embryo development^36,37^. It has been shown that increasing lactate concentration in the culturing media decreases pyruvate uptake rate of preimplantation mouse embryos, suggesting a possible competition between pyruvate and lactate^30^. The kinetic model predicted the steady-state LDH flux with absolute units, which enables the quantification of the fractional contribution of lactate to mitochondrial respiration. We next explore how lactate contributes to the maintenance of mitochondrial flux homeostasis across a wide range of external pyruvate and lactate concentrations.

By fitting the kinetic model to the LDH flux response during an oxygen drop, we predicted the steady state LDH flux for oocytes cultured in AKSOM, 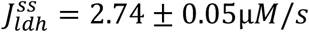. Since LDH converts lactate to pyruvate to power mitochondrial respiration, we asked how much of the respiration rate is accounted for by this LDH flux: i.e. how much of mitochondrial respiration is supported by the uptake of lactate from the media, which is subsequently converted to pyruvate inside the oocyte by LDH (as opposed to pyruvate uptake from the media). To do this, we first convert the LDH flux to the total LDH conversion rate of the oocyte by multiplying 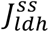 with the cytoplasmic volume of the oocyte *V*_*cyto*_. In this work, we used the relation *V*_*mito*_/*V*_*cyto*_ = 1 from the fluorescence measurement of Mitotracker^19^. By definition, *V*_*oocyte*_ = *V*_*mito*_ + *V*_*cyto*_. Therefore, *V*_*cyto*_ = 1/2*V*_*oocyte*_. *V*_*oocyte*_ = 2 × 10^5^ *μm*^3^ as calculated from the average diameter of a MII oocyte^19^, leading to *V*_*cyto*_ = 1 × 10^5^ *μm*^3^, hence the total conversion rate of LDH (Φ_*ldh*_) in AKSOM is 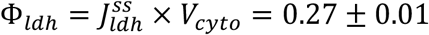 (Figure 6a). The oxygen consumption rate of the mitochondria (Φ_Q_) is calculated from 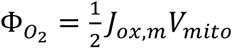, where *J*_*ox,m*_ = 55.4 ± 1.9 *μM*/*s* and *V*_*mito*_ = 1 × 10^5^ *μm*^3^. The factor of 1/2 is to account for the fact that every oxidation of 2NADH consumes 1 O_2_. Therefore, Φ_Q2_ = 2.77 ± 0.09 fmol/s/oocyte (Figure 6a). Given that each lactate contributes to the consumption of 3O_2_, the fraction contribution of lactate to the total oxygen consumption rate of the mitochondria is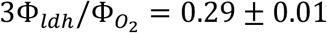. In conclusion, lactate contributes ~30% of the total mitochondrial respiration for MII mouse oocytes cultured in AKSOM.

**Figure 6.**
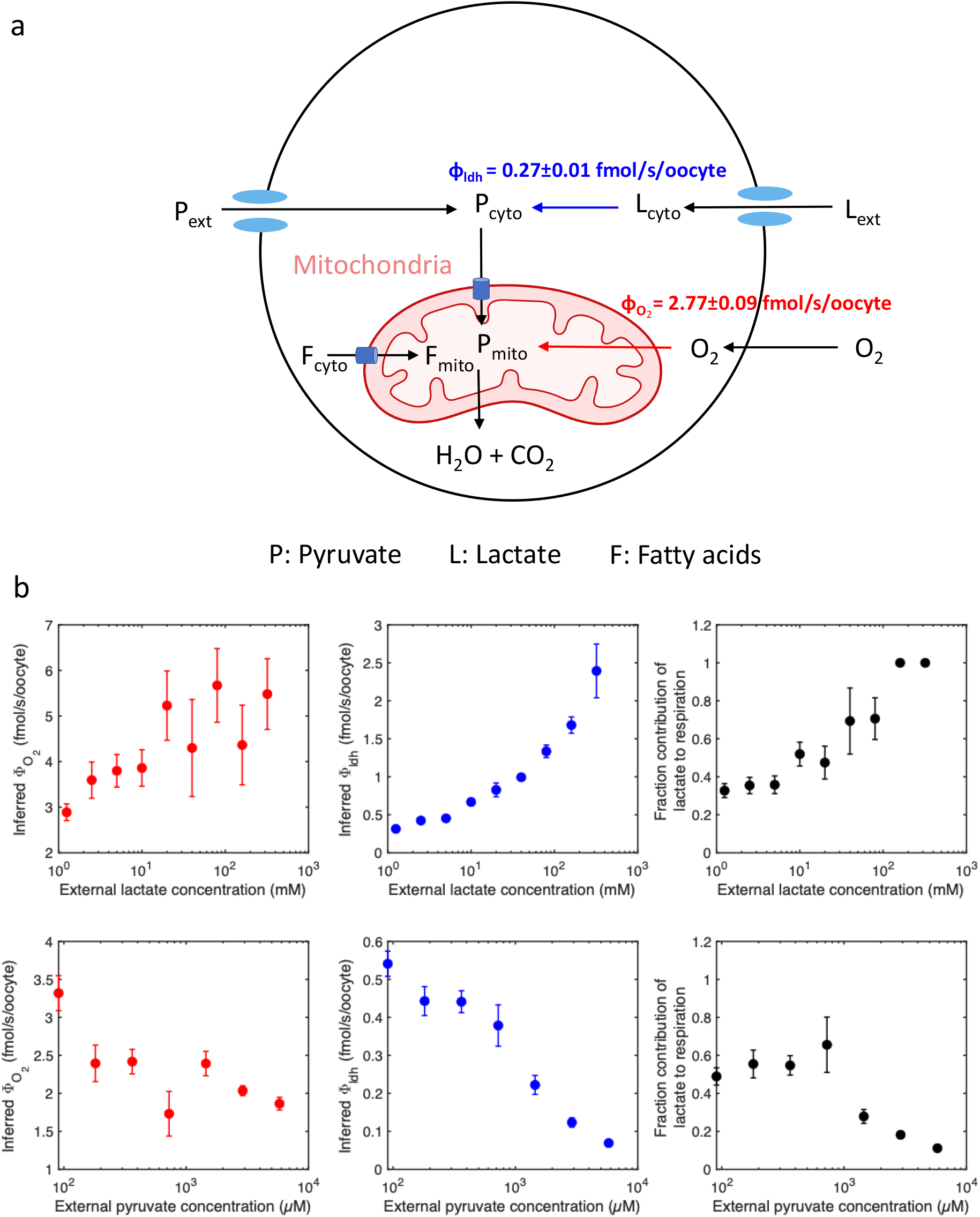
Fractional contribution of lactate to mitochondrial respiration varies significantly while respiration rate remains almost constant in response to nutrient perturbations. a) Schematic showing nutrient flux partitioning in matured MII mouse oocyte in AKSOM media. b) Inferred oocyte oxygen consumption rate 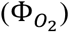, LDH conversion rate (Φ_*ldh*_) and fraction contribution of lactate to mitochondrial respiration for titration of external lactate concentrations with fixed external pyruvate concentration at 181µM (top row) and titration of external pyruvate concentrations with fixed external lactate concentration at 10mM (bottom row). n=18,18,18,18,30,12,12,12,6 for lactate concentrations at 1.25mM, 2.5mM, 5mM, 10mM, 20mM, 40mM, 80mM, 160mM and 320mM, respectively. n=8,7,8,8,8,12,12 for pyruvate concentrations at 90.5µM, 181µM, 362µM, 724µM, 1448µM, 2896µM and 5792µM, respectively. Error bars denote standard error of the mean across individual oocytes. n denotes number of oocytes.

To reveal how much lactate contributes to mitochondrial respiration across different nutrient conditions, we repeated the same procedures for AKSOM in conditions with different external pyruvate and lactate concentrations. First, we obtained *J*_*ldh*_ and *J*_*ox,m*_ using Eqn. (1) from the concentrations of bound and free NADH concentrations at different pyruvate and lactate concentrations (Figure 2c and d). To infer 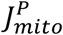, we applied Eqn (1) with a constant *α*_*m*_ = 5.4 ± 0.2*s*^−1^. [*NADH*_*b*_]_*m,eq*_/[*NADH*_*f*_] is obtained at each nutrient condition through oxygen drop experiment and taken to be the lowest value of [*NADH*_*b*_]_*m*_/[*NADH*_*f*_]_*m*_ during the oxygen drop for that specific nutrient condition (Supp Mat Figure 6Sc and d). 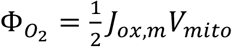 is calculated at each nutrient condition and plotted in Figure 6b left column. To infer *J*_*ldh*_, we used 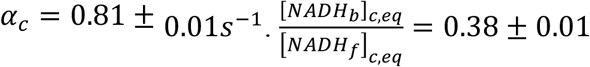 is obtained as the lowest 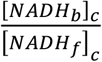 during oxygen drop in the presence of oxamate (Supp Mat Figure 3S b). The same value of 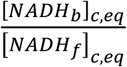 is applied for all nutrient conditions to infer *J*_*ldh*_ (Figure 6b middle column). 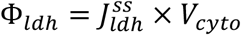 is calculated for all nutrient conditions and plotted in Figure 6b middle column. Finally, we plotted the fraction contribution of lactate to mitochondrial respiration 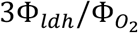 in Figure 6b right column as a function of external pyruvate and lactate concentrations. We observed that the total conversion rate of lactate increases sharply with external lactate concentrations, while the oxygen consumption rate of the oocyte remains almost constant over a wide range of external lactate concentrations. The latter is consistent with mitochondrial flux homeostasis. As a result, the fraction contribution of lactate to mitochondrial respiration rate increases sharply with the external lactate concentrations (Figure 6b first row right). On the other hand, the total conversion rate of lactate decreases with external pyruvate concentrations, but due to the insensitivity of the oxygen consumption rate, the fraction contribution of lactate to mitochondrial respiration rate decreases with external pyruvate concentrations (Figure 6b second row right). As a consistency check, the contribution of lactate to mitochondrial respiration is negligible in the absence of external lactate concentrations (Supp Mat Figure 7S). The pyruvate concentration in the mouse oviduct is estimated to be in the range of 0.14-0.37mM while the lactate concentration is in the range of 4.26-4.79mM^38^. From Figure 6b we see that at external pyruvate concentration of 181µM and lactate concentration of 5mM, which is close to the physiological condition, the predicted fraction contribution of lactate to respiration is 36 ± 5%. Therefore, lactate contributes significantly to mitochondrial respiration in mouse oocytes under physiologically relevant pyruvate and lactate concentrations.

Remarkably, while the total conversion rate of lactate changes by nearly two orders of magnitude across all nutrient conditions (Figure 6b middle), the oxygen consumption rate only varies by a factor of ~3 (Figure 6b left). Our observation suggests that oocytes maintain a homeostatic mitochondrial respiration rate by modulating the contribution of lactate to respiration through the conversion of lactate to pyruvate via lactate dehydrogenase. The fractional contribution of lactate to mitochondrial respiration increases with external lactate concentrations and decreases with external pyruvate concentrations.

## Discussion

In this work, we have observed distinct behaviors of cytoplasmic and mitochondrial NADH levels in response to oxygen depletion and recovery in matured mouse oocytes, where cytoplasmic NADH level displays a transient overshoot upon oxygen recovery. Through specific biochemical perturbations, we have shown that cytoplasmic NADH levels are coupled to mitochondrial NADH levels by lactate dehydrogenase (LDH) and mitochondrial pyruvate uptake fluxes. Inhibiting LDH or mitochondrial pyruvate uptake suppresses the cytoplasmic NADH response to oxygen perturbation. Monitoring LDH activities upon titration of external pyruvate and lactate concentrations reveals that the net reaction of lactate dehydrogenase operates in the direction of converting lactate to pyruvate to support mitochondrial respiration, therefore rendering lactate a nutrient for mouse oocytes. This result is consistent with the fact that LDH-B isoform, which converts lactate to pyruvate, is the most abundant LDH isoform in mouse oocytes^35^. We applied a coarse-grained NADH redox model to predict the dynamics of flux through LDH and mitochondrial pyruvate uptake flux during oxygen drop and recovery. We discovered that LDH flux decreases with decreasing mitochondrial pyruvate uptake flux and overshoots upon a fast recovery of mitochondrial pyruvate uptake flux.

To explain the dynamical coupling between cytoplasmic LDH flux and mitochondrial pyruvate uptake flux, we developed a kinetic model of the dynamics of cytoplasmic pyruvate and lactate in mouse oocytes due to the exchange of pyruvate and lactate with the media, the conversion between lactate and pyruvate by LDH, and the uptake of pyruvate into mitochondria. Linearizing around the steady state before the oxygen perturbation and using the transfer function analysis, we have analytically solved the response of LDH flux to a step increase of mitochondrial pyruvate uptake flux. Our solution indicates that the overshoot of LDH flux occurs as a result of the competition of intrinsic timescales determined by the exchange rates of pyruvate and lactate with the media and their interconversion rates catalyzed by LDH. We confirmed the model prediction through numerical solution of the linearized kinetic model and quantitatively fitted the full dynamical response of LDH flux in response to changes of mitochondrial pyruvate uptake flux during oxygen drop and recovery. The fitting yields quantitative predictions for pyruvate and lactate uptake rates, LDH catalysis rates and the steady-state LDH flux before the oxygen perturbation. Remarkably, after reducing the LDH catalysis rates by 80% with the LDH inhibitor oxamate, the model still fully captures the measured response of LDH flux to the changes of mitochondrial pyruvate uptake flux. Particularly, the model successfully predicts the suppression of the overshoot dynamics of LDH flux by oxamate.

Finally, using the fitted steady-state LDH flux from oocytes in AKSOM, we calibrated the NADH redox model by determining the rate constant *α*_*c*_ for the cytoplasm (Equation 1), which enables the prediction of LDH flux in other conditions. The rate constant *α*_*m*_ for the mitochondria is calibrated from the measurement of oxygen consumption rate of the oocytes^19^. Using the calibrated NADH redox model, we predicted the absolute values of oxygen consumption rate and LDH flux integrated over a single oocyte for the titration of external pyruvate and lactate concentrations. The model predicts that lactate contributes to a significant fraction of mitochondrial respiration over a wide range of external pyruvate and lactate concentrations. The lactate contribution increases with external lactate concentration and decreases with external pyruvate concentration. At external pyruvate concentration of 181µM and lactate concentration of 5mM, which is close to the physiological condition, our model predicted a fraction contribution of lactate to respiration of 36 ± 5%. This value is also close to the AKSOM condition, which is optimized for embryo development, where our model predicts that lactate contributes to 29 ± 1% of mitochondrial respiration. We further discovered that while the fractional contribution of lactate to mitochondrial respiration varies drastically in response to changes of external lactate and pyruvate concentrations, mitochondrial respiration rate is remarkably insensitive to these nutrient perturbations. This is consistent with the observation of flux homeostasis in earlier works^14,19^. Our work suggests that the oocytes maintain a homeostatic mitochondrial respiration rate by modulating the contribution of lactate to mitochondrial respiration.

We note that we have inferred the LDH flux from cytoplasmic NADH concentrations using the coarse-grained NADH redox, which assumes that NADH redox cycle is at a quasi-steady state in the cytoplasm despite the presence of the transient NADH overshoot dynamics. This assumption holds when *J*_*ox,c*_ is much larger than the rate of change of [*NADH*_*f*_]_*c*_. From Figure 5e, we inferred that *J*_*ox,c*_~180*μM*/*min* during the overshoot. From Figure 4b, we estimated that the rate of change of [*NADH*_*f*_]_*c*_ is ~1.5*μM*/*min*. Hence the quasi-steady state assumption holds for cytoplasmic NADH cycle even during the overshoot.

To infer NADH oxidation fluxes using eqn 1, the equilibrium NADH ratio and the rate constants need to be obtained for both the mitochondria and the cytoplasm. The equilibrium NADH bound ratio for the mitochondria is obtained at the lowest oxygen level, and obtained at each perturbation conditions, including oxamate perturbation (Supp Mat Figure 3Sa) and nutrient titration (Supp Mat Figure 6Sc-d) through oxygen drop. The equilibrium NADH bound ratio displays moderate variations with respect to nutrient perturbations and is hence accounted for by using the corresponding value at each condition. However, the equilibrium NADH bound ratio in the cytoplasm is chosen to be a constant obtained as the lowest cytoplasmic NADH bound during the oxygen drop in the presence of sodium oxamate (Supp Mat Figure 3Sb). This is a choice for simplicity and more work needs to be done to test the robustness of the equilibrium cytoplasmic NADH bound ratio in the future. Furthermore, we also assumed that the rate constants in Eqn 1 to be insensitive to all perturbations considered so far. These constants are obtained in the AKSOM condition and generalized to all other conditions. The constancy of the mitochondrial rate parameter has been verified by the fact that the NADH redox model quantitatively predicts the OCR across a wide range of perturbations^19^. The constancy of the cytoplasmic rate parameter, however, remains to be tested through direct measurement of LDH flux in the future.

It has long been a technical challenge to measure intracellular metabolic fluxes with subcellular resolution in living cells. Metabolic flux analysis (MFA) has been widely used to infer intracellular metabolic fluxes from isotope labeling of tracer metabolites using mass spectrometry^5^, but this method requires the flash freezing of the sample and is not performed on live cell. Fluorescent biosensors have been developed to measure levels of intracellular metabolites such as ATP^39^, glucose^40^, pyruvate^41^ and lactate^42^, but they do not directly provide readout of the turnover rates of these metabolites. Mitochondrial pyruvate uptake rate has been inferred by measuring the rate of depletion of cytoplasmic pyruvate level using a fluorescent pyruvate biosensor after inhibition of cellular pyruvate transporter^43^, but this interpretation requires acute metabolic perturbation and the absence of other pyruvate producing or consuming pathways. We have demonstrated in this work that we can apply the NADH redox model to infer the NADH oxidation fluxes in both mitochondria and cytoplasm in intact living cells. The inferred NADH oxidation flux in the mitochondria corresponds to the ETC flux, which has been verified through direct oxygen consumption rate measurements of the oocytes^19^. The inferred NADH oxidation flux in the cytoplasm equals the conversion flux of lactate to pyruvate through lactate dehydrogenase in the oocytes. The predicted LDH flux responds to lactate and pyruvate titrations in a self-consistent manner, and the dynamical response of the inferred LDH flux can be explained quantitatively by the kinetic model. To test the accuracy of the inferred LDH flux, a direct measurement of the LDH flux is required. Stimulated Raman Spectroscopy could provide a potential technique to measure nutrient uptake rates with single cell resolution in living cells^44^, and could be a test for our model predictions in the future.

While dynamical control theory has been applied to study the sensitivity and robustness of metabolic networks from a theoretical perspective^45^, it has rarely been applied to analyze biological data. The transfer function analysis used in this work provides an example of how to implement dynamical control theory to analyze experimental data to gain information about metabolic kinetics from transient flux dynamics in biological systems. Control theory provides analytical solutions for the response of LDH flux (*j*_*ldh*_(*t*)) to the step change and linear ramp of the mitochondrial pyruvate uptake flux 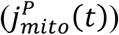. The theory predicts an overshoot dynamics of *j*_*ldh*_(*t*) in response to a step change in 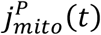 (Supp Mat Eqn S23) and a delayed response to a linear ramp of 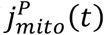 (Supp Mat Eqn S29). Both behaviors have been observed experimentally (Figure 4b). We have shown that the inhibition of lactate dehydrogenase diminishes the overshoot of *j*_*ldh*_(*t*) by reducing the ratio between the characteristic timescales of the transfer function. Our work demonstrates the power of using control theory to explain complex temporal dynamics of metabolic fluxes. Transient overshoot of NAD(P)H signals has also been observed in neurons in response to depolarizing pulses. This overshoot has been attributed to calcium entrance via mitochondrial calcium uniporter that stimulates TCA cycle^15^. We expect our control theory analysis cold also be applicable to neuron stimulation, and could thus potentially reveal relevant kinetic rates. Glycolytic oscillation is another well-known metabolic dynamic where NADH levels are observed to undergo damped or sustained oscillations in response to metabolic perturbations such as cyanide poisoning or glucose feeding after starvation^46^. Control theory has been applied to model this behavior, which proposes that glycolytic oscillation is an inevitable side effect due to trade-offs between robustness and efficiency involving feedback control^47^. Our current kinetic model does not involve any feedback controls and is built entirely on the law of mass action. We have thus demonstrated that simple laws of mass action can explain seemingly complex dynamical responses in cells. For more complicated dynamical cellular responses, feedback control modules can be added to the kinetic model, and the transfer function formulism provides a powerful tool to analyze the dynamical responses by treating the cell as a dynamical system as in engineering systems^7^.

Finally, we expect our NADH redox model and the control theory approach to be a powerful tool in dissecting the dynamics of metabolic pathways in other living cells. They provide a method to infer intrinsic rates and timescales of metabolic pathways subject to transient metabolic perturbations with the potential of revealing new metabolic control mechanisms.

## Methods and Materials

### Oocytes culturing and oxygen perturbation

Frozen MII mouse oocytes (Strain B6C3F1) were purchased from EmbryoTech. Oocytes were thawed and cultured in droplets of AKSOM media (MilliporeSigma) in plastic petri dish. Mineral oil (VitroLife) was applied to cover the droplets to prevent evaporation of the media. The plastic dish was placed in the incubator at 37C, 5% CO2 and 20% O2 before imaging. For imaging, oocytes were cultured in a 2uL AKSOM media droplet and covered with 400-500ul of oil in a 35mm glass bottom confocal dish (FluoroDish, WPI). The dish was placed in an ibidi chamber at 37C and 5% CO_2_ during imaging. For oxygen perturbation, two gas tanks with 5% O_2_ and 0% O_2,_ respectively, were connected to a single tube through a Y-splitter. The tube was connected to the ibidi chamber. Pressures of both tanks are manually tuned gradually to drop oxygen continuously from 5% to 0% in the chamber. The oxygen level in the chamber is measured using the oxygen sensor from gaslab (model: CM-42991).

### Nutrient and drug perturbations

Nutrient perturbations are achieved by making KSOM media following Cold Spring Harbor Laboratory protocols and titrating pyruvate and lactate concentrations while keeping all other media compositions constant as in the standard KSOM media. For pyruvate titration, lactate level is maintained at 10mM. For lactate titration, pyruvate level is maintained at 181uM. Drug perturbations are achieved by adding inhibitors to AKSOM media obtained from MilliporeSigma.

### Fluorescence lifetime imaging of NADH

FLIM of NADH was performed using a two-photon scanning confocal microscope with a 40X 1.25NA water immersion Nikon objective, Becker and Hickle Time Correlated Single Photon Counting (TCSPC) acquisition system with a hybrid detector (HPM-100-40) and a pulsed Insight X3 femtosecond laser (Spectra-Physics). NADH autofluorescence was excited at 750nm and collected with a 460/50 nm emission filter. Laser power was maintained at 3mW at the objective during imaging. 512 by 512 pixel images were collected with a pixel size of 420nm and 30s acquisition time per frame. The pixel dwell time is 4.09us.

### Two-exponential fitting of NADH decay curve

We fitted the NADH decay curve with *G*(*τ*) = *IRF*(*τ*) ∗ [*C*_1_*F*(*τ*) + *C*_2_]. *IRF*(*τ*) is the instrument response function collected using a urea crystal. 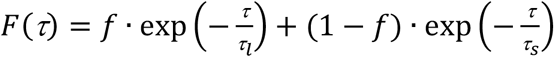 is the two-exponential model, where *τ*_*l*_ and *τ*_*s*_ are the long and short fluorescence lifetimes and *f* is the fraction of bound NADH. *C*_1_ is the amplitude of the decay and *C*_2_ is the background noise. We implemented least-square fitting using the Levenberg-Marquardt algorithm with a custom-built Matlab code.

### Calculation of free and bound NADH concentrations

The concentrations of free ([NADH_f_]) and ([NADH_b_]) are obtained using the relation 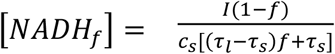 and 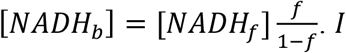 is the NADH fluorescence intensity obtained by integrating the decay curve *G*(*τ*) and divide by the number of pixels. *c*_*s*_ is a calibration factor that is obtained through in vitro calibration of the NADH solutions^19^.

### Inference of NADH oxidation flux

NADH oxidation flux is defined as the number of NADH molecules oxidized per unit of time and per unit of volume, which has the unit of concentration per time. Using a coarse-grained NADH redox model^19^, we derived the NADH oxidation flux at quasi-steady state as 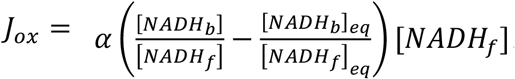 is a rate constant. [*NADH*_*b*_]_*eq*_ and [*NADH*_*f*_]_*eq*_ are bound and free NADH concentrations at equilibrium, where *J*_*ox*_ = 0. In mitochondria, *J*_*ox,m*_ is proportional to the oxygen consumption rate (OCR) of mitochondria. From a direct measurement of OCR, 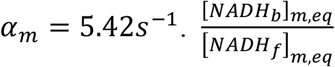 in the mitochondria is obtained as the bound and free NADH ratio at the lowest oxygen level (Supp Mat Figure 2S d). In the cytoplasm at quasi-steady state, *J*_*ox,c*_ is proportional to the LDH flux. Inferring steady-state LDH flux from the kinetic model provides a calibration for *α*_*c*_ in the cytoplasm, which yields 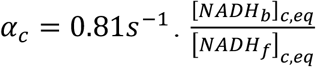 in the cytoplasm is obtained with 10mM of sodium oxamate in the media and at the lowest oxygen level (Supp Mat Figure 3S b).

### Oxygen consumption rate measurements

Oxygen consumption rate (OCR) was measured using a nanorespirometer system from Unisense^48^. 10 oocytes were placed at the bottom of a thin capillary well 0.68mm in diameter and 3mm in depth. The top of the well is open and exposed to atmospheric oxygen level, the rest of the well is sealed. The well was immersed in AKSOM media. As the oocytes consume oxygen at the bottom of the well, a linear oxygen gradient was established within the capillary well at steady state. A motor-controlled oxygen-sensing electrode was used to measure this oxygen gradient. The OCR was computed by multiplying the slope of the oxygen gradient with the diffusivity of oxygen and cross-sectional area of the well. The OCR per oocyte is computed by dividing the total OCR by the number of oocytes.

### Cellular pyruvate uptake rate measurements

Cellular pyruvate uptake rate was obtained by measuring the decrease of the pyruvate concentration in the spent media using the Pyruvate Assay Kit (Cayman 700470). 20 oocytes were cultured in 2ul droplet of media with 0.02g/l pyruvate and no lactate for 5 hours at 37C and 20% O_2_. 2uL of spent media was pipetted out and diluted in 18ul of buffer media from the pyruvate assay to make 20ul sample. The sample was then mixed in a 96 plate well with fluorescence detector, cofactor and enzyme mixtures from the pyruvate assay to initiate fluorescence chemical reactions. The fluorescence signal, which is proportional to the concentration of pyruvate in the sample, was detected using the Neo2 plate reader with 530/10nm excitation and 580/10nm emission. The fluorescence signal was calibrated with a standard pyruvate titration to yield absolute concentrations of pyruvate in the sample. The pyruvate uptake rate per oocyte was calculated as the rate of decrease of pyruvate concentration in the spent media by dividing the change of pyruvate concentration in the media by the culturing time of the oocytes and the number of oocytes.

### Calculation of mitochondrial pyruvate uptake flux

Mitochondrial pyruvate uptake flux is calculated using the equation 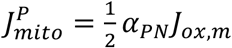, where the ½ factor accounts for the observation that mitochondrial pyruvate uptake accounts for half of the mitochondrial OCR. 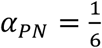 is the stoichiometry factor, resulting from the fact that the NADH to O_2_ stoichiometry is 2 to 1 and O_2_ to pyruvate stoichiometry is 3 to 1 if pyruvate is converted from lactate. If pyruvate is taken up from the media, the O_2_ to pyruvate stoichiometry is 2.5 to 1 because of the lack of cytoplasmic NADH to be transferred into the mitochondria. For simplicity, we consistently choose O_2_ to pyruvate stoichiometry to be 3 to 1.

### Numerical solution of the kinetic model and model parameter fitting

The linearized kinetic model as defined in Eqn S7 and S8 is solved numerically with an initial guess of the numerical values of the kinetic parameters *k*_*l*_ = 0.002*s*^−1^, *k*_*p*_ = 0.4*s*^−1^, *r*_*l*_ = 0.01*s*^−1^ and *r*_*p*_ = 0.55*s*^−1^. The time-dependent 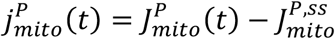 during oxygen drop and recovery and *p*(0) = 0 and *l*(0) = 0 are provided as the input. For numerical stability, 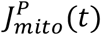 is smoothened by fitting piece-wise linear functions to different regions of 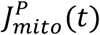. The cytoplasmic pyruvate concentration deviation *p*(*t*) and lactate concentration deviation *l*(*t*) during oxygen drop and recovery is obtained by solving eqn 4 and 5 numerically using finite difference scheme. The LDH flux deviation from steady-state 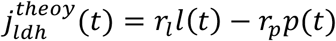 is computed as the output. The full LDH flux from the numerical solution is 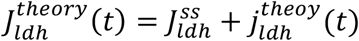. The kinetic parameters and 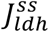 are adjusted by fitting the simulated 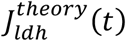 to the experimentally measured 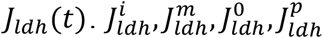 and 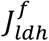 denote the initial steady-state *J*_*ldh*_ before oxygen drop, at the middle point of the oxygen drop, at the lowest oxygen level, at the maximum value of *J*_*ldh*_after the oxygen recovery, and at the restored steady-state after oxygen recovery, respectively. *τ*_*relax*_ denotes the relaxation time scale of *J*_*ldh*_ going from 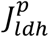 to 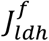 fitted with an exponential decay. The simulated *J*_*ldh*_(*t*) is fitted to match the experimentally measured *J*_*ldh*_(*t*) by fitting 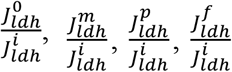 and *τ*_*relax*_ using least-square fitting Levenberg-Marquardt algorithm. The best-fitting kinetic parameters are *k*_*l*_ = 0.0028 ± 0.0006, *k*_*p*_ = 0.3393 ± 0.1261*s*^−1^, *r*_*l*_ = 0.0086±0.0041*s*^−1^, *r*_*p*_ = 0.6927 ± 0.0637*s*^−1^ and 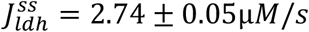 at the AKSOM condition. We performed the same procedure for the oxamate condition, except that we fixed 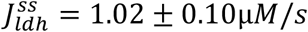, *k*_*l*_ = 0.0028*s*^−1^, *k*_*p*_ = 0.3393*s*^−1^, and only fitted a multiplicative factor *λ* that scales 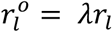 and 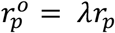 by fitting 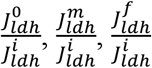, which yields *λ* = 0.20 ± 0.01.

## Acknowledgement

We thank Mike van der Naald for his critical feedback on the manuscript especially his insights on the interpretation of the timescales governing the overshoot dynamics. We thank Efe Ilker and Alphée Michelot for their feedback on the abstract. This study received support from the Deutsche Forschungsgemeinschaft (DFG; German Research Foundation) under Germany’s Excellence Strategy — EXC-2068-390729961 — Cluster of Excellence Physics of Life of TU Dresden. This work is supported by the National Institutes of Health (R01HD092550-01) and the National Science Foundation (PFI-TT-1827309, PHY-2013874, and MCB-2052305).

## Author contributions

X.Y. and D.J.N. designed research; X.Y. performed research; X.Y. and D.J.N. developed theoretical models; X.Y. analyzed data; X.Y. and D.J.N. wrote the paper.

## Supplementary Materials

### Kinetic Model for Pyruvate and Lactate Metabolism

We developed a kinetic model that governs the dynamics of cytoplasmic pyruvate and lactate concentrations (*P*_*cyto*_ and *L*_*cyto*_) by accounting for pyruvate and lactate uptake from the media (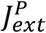 and 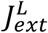), mitochondrial pyruvate uptake 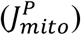 and lactate dehydrogenase flux (*J*_*ldh*_).

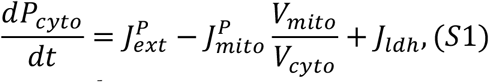

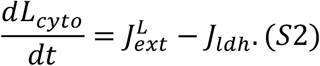

where 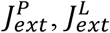 and *J*_*ldh*_ are generally non-linear functions of cytoplasmic pyruvate and lactate concentrations (*P*_*cyto*_, *L*_*cyto*_) and external pyruvate and lactate concentrations (*P*_*ext*_,*L*_*ext*_). We consider perturbations around steady state 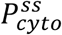, 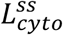:

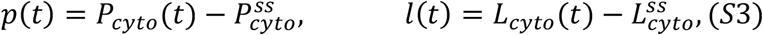

And define deviation of fluxes from steady-state values:

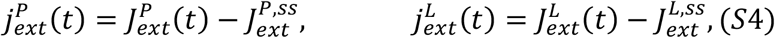

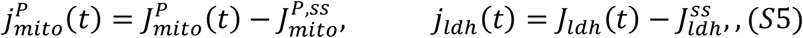

Considering perturbations that lead to changes in cytoplasmic pyruvate and lactate concentrations but keep external pyruvate and lactate concentrations constant, we can expand 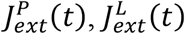 and *J*_*ldh*_(*t*) up to the first order in *p*(*t*) and *l*(*t*) and obtain the linearlized equations for the dynamics of *p*(*t*) and *l*(*t*):

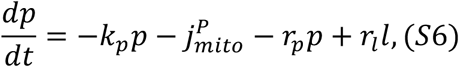

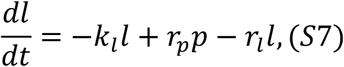

where

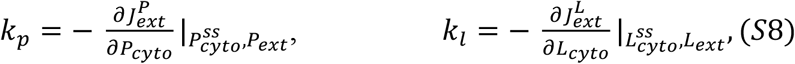

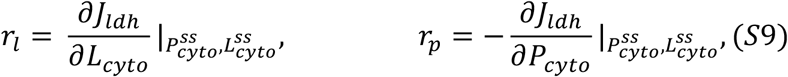

and 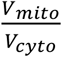 from mitotracker measurement^1^.

Our goal is to predict the dynamics of the flux through LDH (*j*_*ldh*_(*t*) = *r*_*l*_*l*(*t*) − *r*_*p*_*p*(*t*)) in response to the change of mitochondrial pyruvate consumption flux 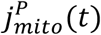.

From Eqn (S6) and (S7), we obtain the second order dynamical equations for *p*(*t*)

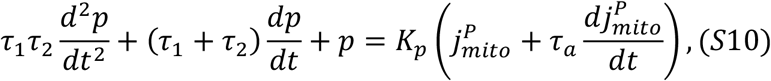

and *l*(*t*)

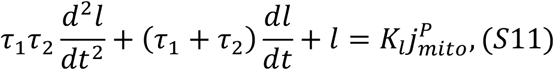

where

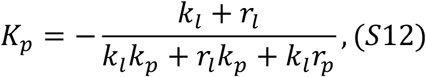

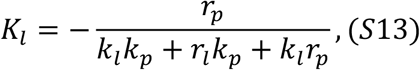

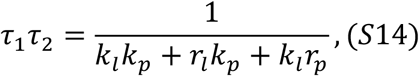

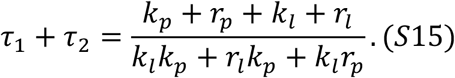

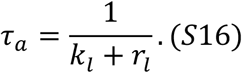

Next, we combined equations (S10) and (S11) to get the dynamical equation for flux through LDH *j*_*ldh*_(*t*) = *r*_*l*_*l*(*t*) − *r*_*p*_*p*(*t*):

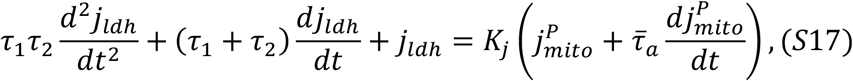

where

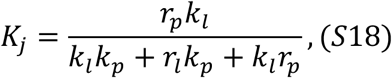

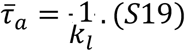

Notice equation (S17) describes a damped oscillator with a driving force. *τ*_1_ and *τ*_2_ are the intrinsic timescales of the oscillator, while 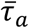 is the timescale governing the response of the oscillator to time-varying perturbations from the driving force.

### Transfer function connecting *j*_*ldh*_(*t*) to 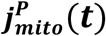

Our goal is to predict the response of LDH flux *j*_*ldh*_(*t*) to time-varying perturbations of the input variable 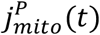. Transfer function in control theory serves this goal. Performing Laplace transform of equation (S17), we obtained the transfer function in the frequency space connecting *j*_*ldh*_(*t*) to 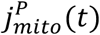

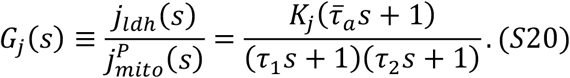

#### Response of *j*_*ldh*_(*t*) to step change in 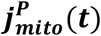

Now let’s consider the response of *j*_*ldh*_(*t*) to a step change in 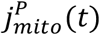, which approximates the regime of sudden oxygen recovery in our experiment.

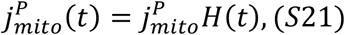

where *H*(*t*) is the Heaviside step function. Laplace transform yields

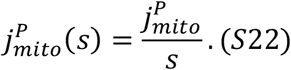

Substituting equation (S22) to (S20) and performing inverse Laplace transform yields

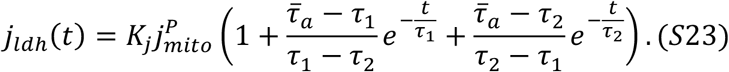

Response to step change as described by equation (S23) has been well characterized in control theory. First, oscillation is absent in our system because *τ*_1_ and *τ*_2_ are always real numbers regardless of the model parameters, i.e. we are in the overdamped limit. In this limit, it is well-known that overshoot of *j*_*ldh*_(*t*) occurs when^2^

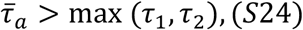

In the limit *r*_*p*_, *k*_*p*_ ≫ *r*_*l*_, *k*_*l*_, which is consistent with the values of the fitted parameters *k*_*l*_ = 0.0028 ± 0.0006*s*^−1^, *k*_*p*_ = 0.3 ± 0.1*s*^−1^, *r*_*l*_ = 0.0086±0.004*s*^−1^ and *r*_*p*_ = 0.69 ± 0.06*s*^−1^ from the main text and referring to equation (S14)-(S15) and (S19), we get

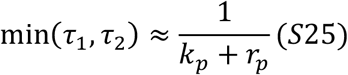

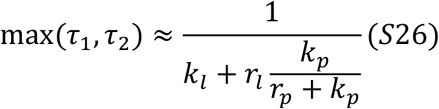

Using the fitted parameter values, we get min(*τ*_1_, *τ*_2_) = 1.01 ± 0.12*s* and max(*τ*_1_, *τ*_2_) = 184.98 ± 50.99*s*. According to Eqn (S23), the longer timescale determines the relaxation time of the LDH flux after oxygen recovery. Inserting Eqn (S19) and (S26) into Eqn (S24), the overshoot condition can be written as

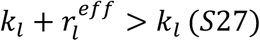

where

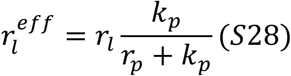

#### Response of *j*_*ldh*_(*t*) to linear decrease of 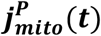

During the oxygen drop, the change of 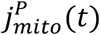 can be approximated as a linear ramp.

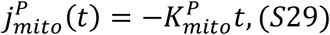

where 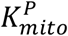 is the rate of change of 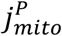. Laplace transform yields

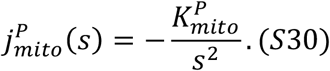

Substituting equation (S30) into (S20) and performing inverse Laplace transform yields

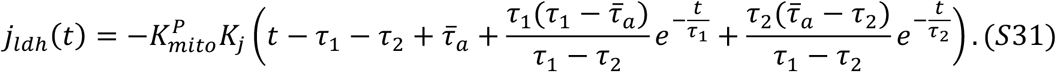

In the limit *t* ≫ *τ*_1_ and *t* ≫ *τ*_2_, equation (S31) reduces to

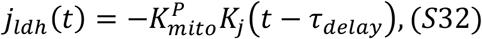

Where 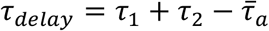. Notice that the response of *j*_*ldh*_(*t*) to a linear ramp of 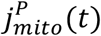 is characterized by a linear response with a time delay.

### Transfer function connecting NADH concentration to 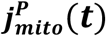

In our experiment, we are directly measuring the concentrations of NADH. The kinetic model for pyruvate and lactate does not explicitly account for the dynamics of NADH, where the NADH dynamics is implied in the LDH flux. Particularly, NADH concentration is assumed to be proportional to LDH flux. This assumption holds in the quasistatic limit where the dynamics of NADH is faster than that of pyruvate and lactate. This assumption is reasonable if, for example, the transport of pyruvate and lactate through the cell membrane is slower than the bulk reactions inside the cell. Given the large volume of the oocyte, this could be the case. This assumption does not have to hold generally, hence in this section we consider a kinetic model where the dynamics of cytoplasmic NADH is explicitly modeled. In addition to the kinetic model of pyruvate and lactate, we model the cytoplasmic NADH dynamics as

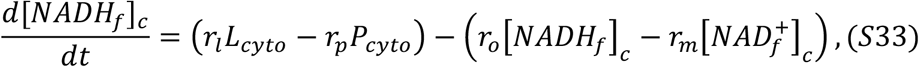

Where *r*_*o*_ and *r*_*m*_ are the forward and reverse reaction rates of malate dehydrogenase that oxidizes NADH in the cytoplasm. Assuming the sum of NADH and NAD+ concentration is a constant

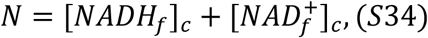

and linearly expand around the steady state of NADH

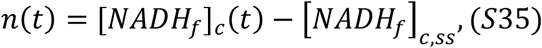

We obtained the linearized equations for the kinetic model

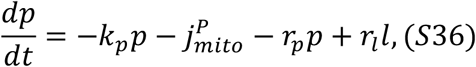

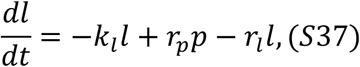

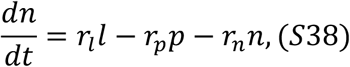

where

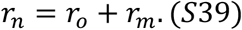

The transfer function connecting *n*(*s*) to 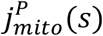 is

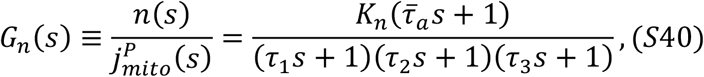

where

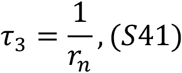

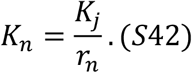

Inverse Laplace transform yields the response of *n*(*t*) to step change in 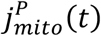

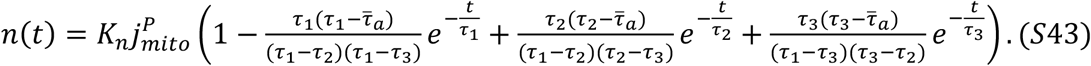

The overshoot in *n*(*t*) occurs when

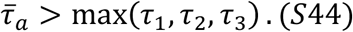

Notice the kinetic model involving NADH dynamics reduces to the kinetic model without NADH dynamics when

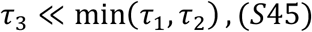

i.e. when the dynamics of NADH is faster than that of pyruvate and lactate.

**Figure 1S.**
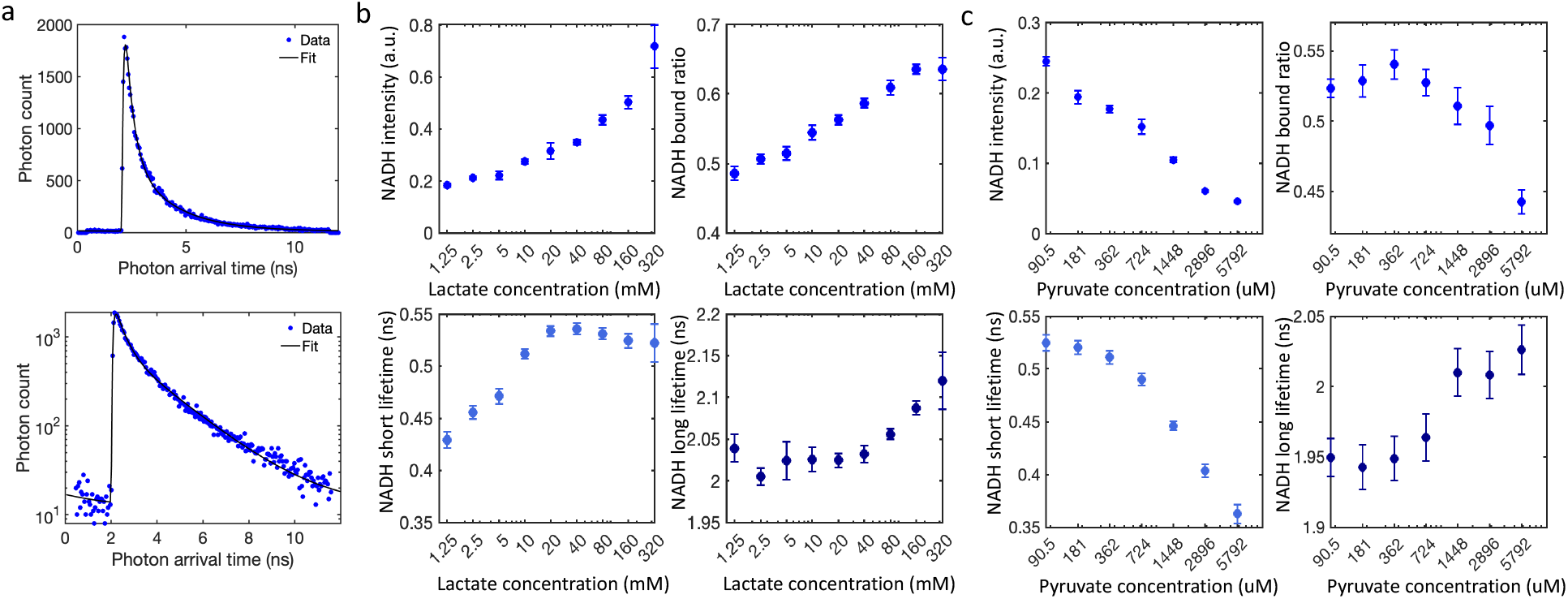
FLIM measurements of cytoplasmic NADH under lactate and pyruvate titrations. a) FLIM curve and the double exponential fitting for cytoplasmic NADH FLIM signal. b) NADH intensity, bound ratio, short and long fluorescence lifetime from double exponential fitting of the FLIM curve as a function of extracellular lactate concentration. n=18,18,18,18,30,12,12,12,6 for lactate concentration from 1.25mM to 320mM, respectively. c) NADH intensity, bound ratio, short and long fluorescence lifetime from double exponential fitting of the FLIM curve as a function of extracellular pyruvate concentration. n=8,7,8,8,8,12,12 for pyruvate concentration from 90.5µM to 5792µM, respectively.

**Figure 2S.**
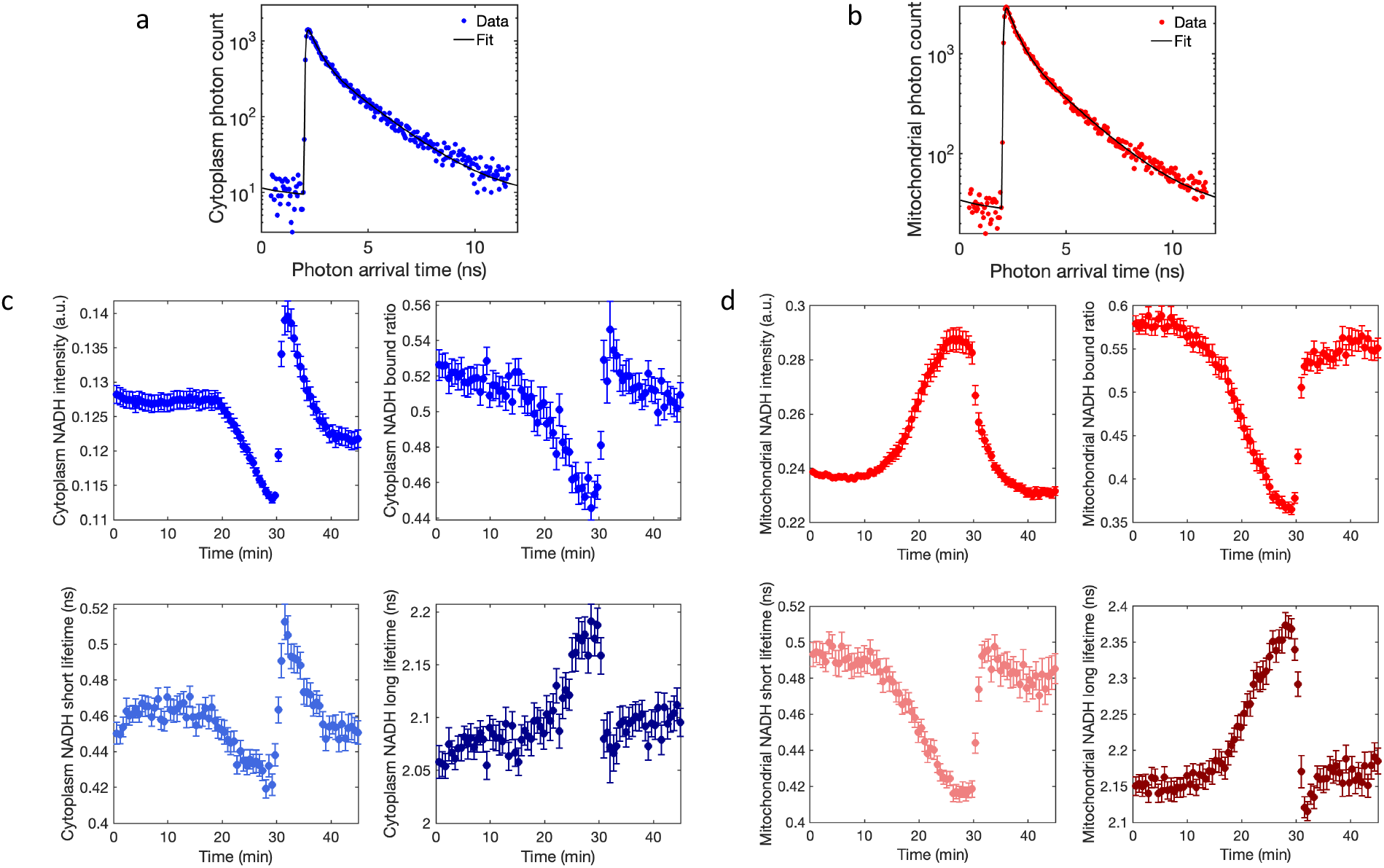
FLIM measurements of mitochondrial and cytoplasmic NADH during oxygen perturbation. a) FLIM curve and double exponential fitting of cytoplasmic NADH. b) FLIM curve and double exponential fitting of mitochondrial NADH. c) Cytoplasmic NADH intensity, bound ratio, short and long fluorescence lifetimes from double exponential fitting during oxygen drop perturbation. d) Mitochondrial NADH intensity, bound ratio, short and long fluorescence lifetimes from double exponential fitting during oxygen drop perturbation. n=68. Error bars denote standard error of the mean across individual oocytes. n denotes number of oocytes.

**Figure 3S.**
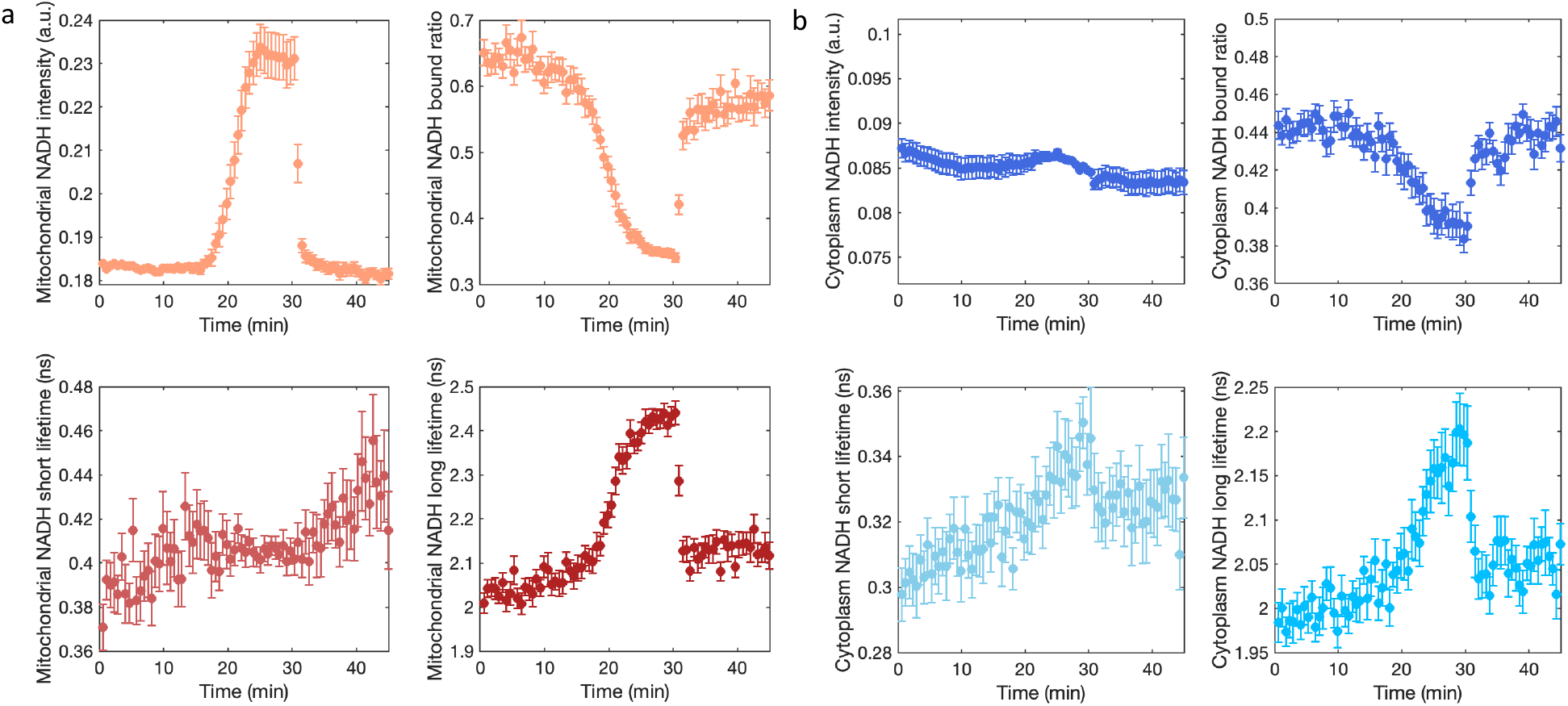
FLIM measurements of mitochondrial and cytoplasmic NADH during oxygen perturbation in the presence of oxamate. a) Mitochondrial NADH intensity, bound ratio, short and long fluorescence lifetime as a function of time during oxygen perturbation. b) Cytoplasmic NADH intensity, bound ratio, short and long fluorescence lifetime as a function of time during oxygen perturbation. n=20. Error bars denote standard error of the mean across individual oocytes. n denotes number of oocytes.

### Oxygen consumption rate under nutrient perturbations

**Figure 4S.**
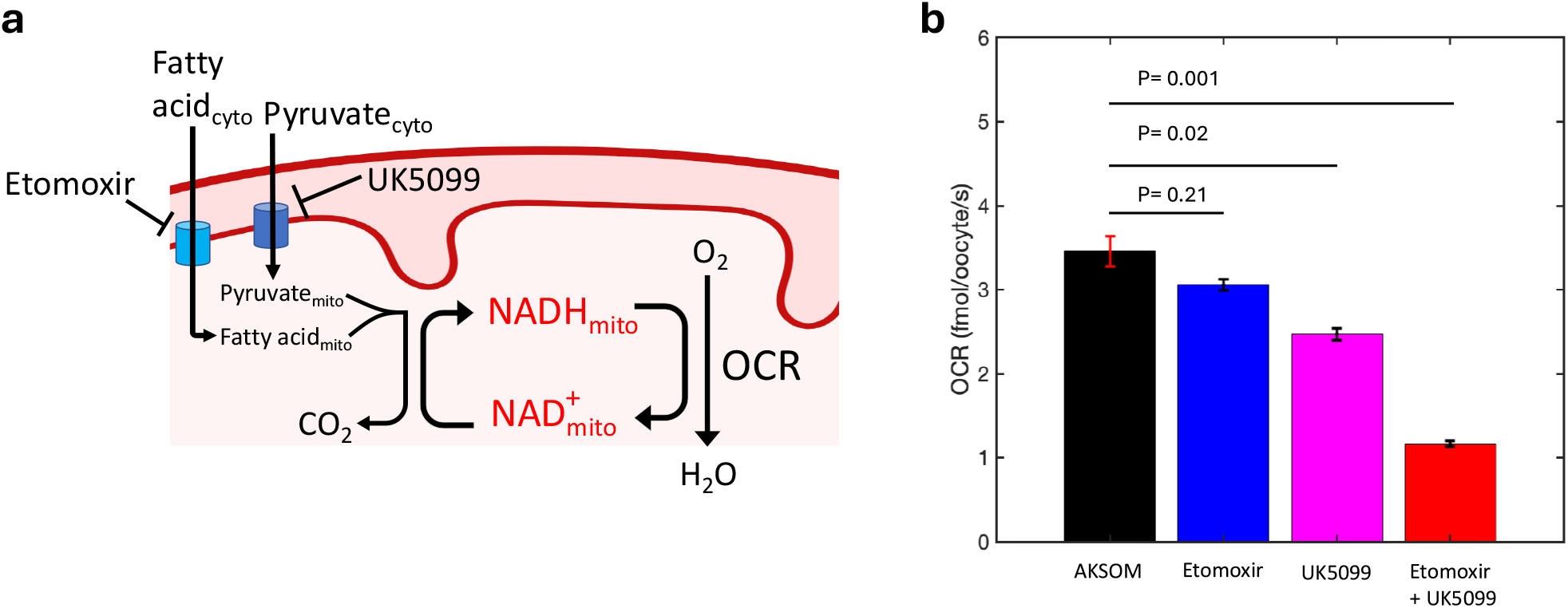
Oxygen consumption rate under nutrient perturbations. a) Schematic of mitochondrial respiration and the inhibitors targeting fatty acid oxidation (etomoxir) and mitochondrial pyruvate uptake (UK5099). b) Oxygen consumption rate measured in different conditions. N=4, 2, 2, 2 for AKSOM, Etomoxir, UK5099 and Etomoxir+UK5099 conditions, respectively. Error bars denote standard error of the mean across different batches of experiment. N denotes number of batch of experiments. Two-sample t-test is performed for statistical testing.

### LDH flux overshoot results from the asymmetric response of cytoplasmic pyruvate and lactate concentrations

**Figure 5S.**
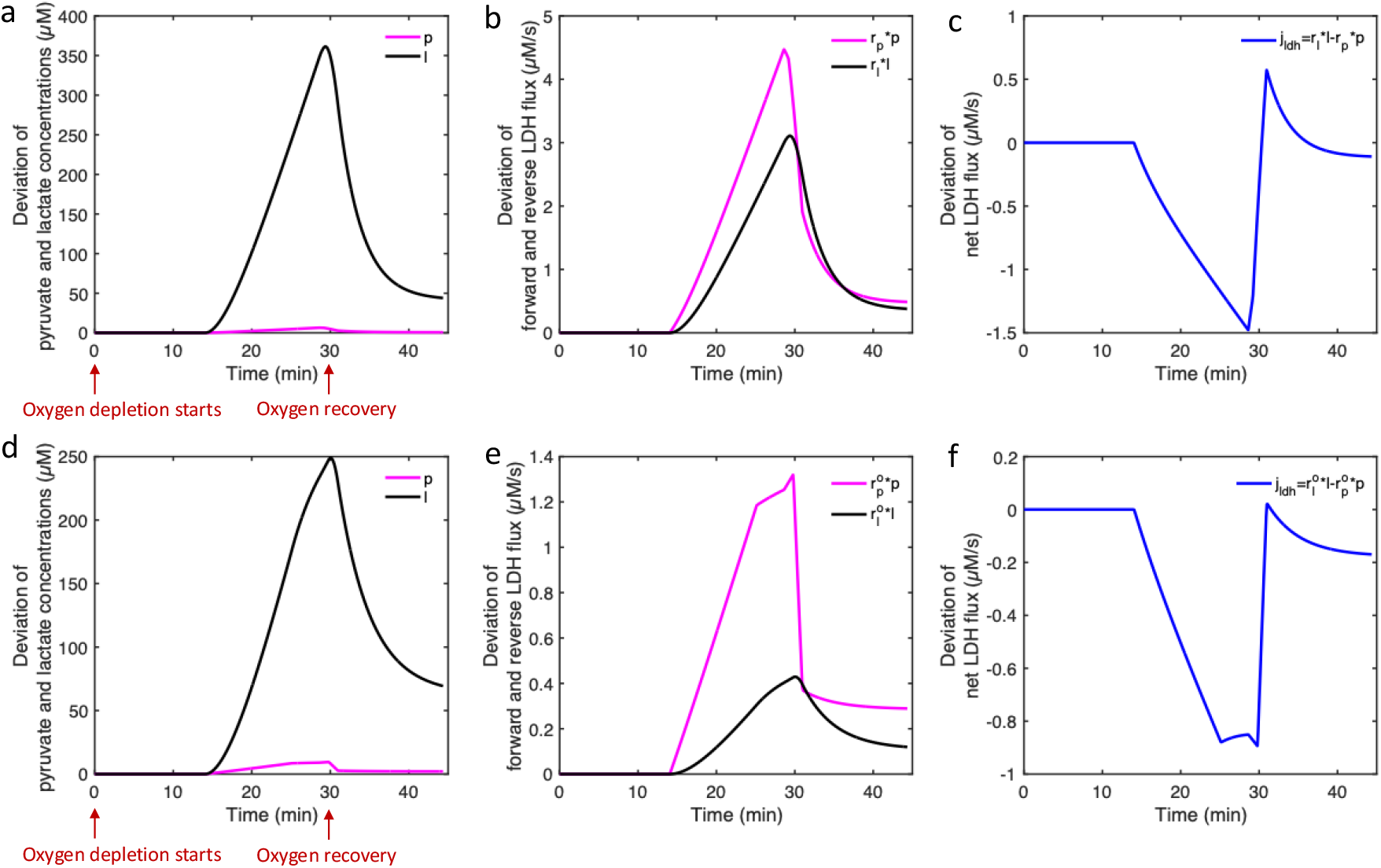
LDH flux overshoot results from the asymmetric response of cytoplasmic pyruvate and lactate concentrations to mitochondrial pyruvate uptake perturbation predicted by the kinetic model. Deviation of cytoplasmic pyruvate and lactate concentrations from initial steady state value for AKSOM (a) and oxamate (b) conditions. Deviation of forward (*r*_*l*_*l*) and reverse (*r*_*p*_*p*) flux of LDH from the initial steady state value for AKSOM (b) and oxamate (e) conditions. Deviation of net LDH flux from the initial steady state value for AKSOM (c) and oxamate (f) conditions.

Control theory has analytically predicted the occurrence of the LDH flux overshoot from the competition of intrinsic timescales determined by the conversion and uptake rates of pyruvate and lactate (Figure 5, main text). To develop an intuitive understanding of the overshoot dynamics, we numerically solved the linearized Eqn S7 and S8 to study how the dynamics of pyruvate and lactate concentrations contribute to the LDH flux overshoot explicitly. Taking the time trace of the deviated mitochondrial pyruvate uptake flux 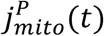 as the input, we predicted the deviated pyruvate and lactate concentrations from their initial steady state values as function of time in the AKSOM condition (Figure 5S a). Multiplying the deviated pyruvate and lactate concentrations by the fitted kinetic parameters *r*_*p*_ = 0.6927 ± 0.0637*s*^−1^ and *r*_*l*_ = 0.0086±0.0041*s*^−1^, respectively, we obtained the deviated forward *r*_*l*_*l* and reverse *r*_*p*_*p* flux of LDH (Figure 5S b). Finally, subtracting the reverse flux from the forward flux, we obtained the net deviated LDH flux throughout time (Figure 5S c). Notice that the overshoot, characterized by a positive value in the deviated net LDH flux, occurs when the deviated forward flux is larger than the reverse flux, which could result from the asymmetry in the decay rate of the forward and reverse LDH flux (Figure 5S b).

We performed the same procedure for the oxamate condition (Figure 5S d-f). Notice that in this case, the forward flux is almost always smaller than the reverse flux (Figure 5S e), hence overshoot does not occur in the deviated net flux (Figure 5S f).

### Mitochondrial NADH concentrations during pyruvate and lactate titration

**Figure 6S.**
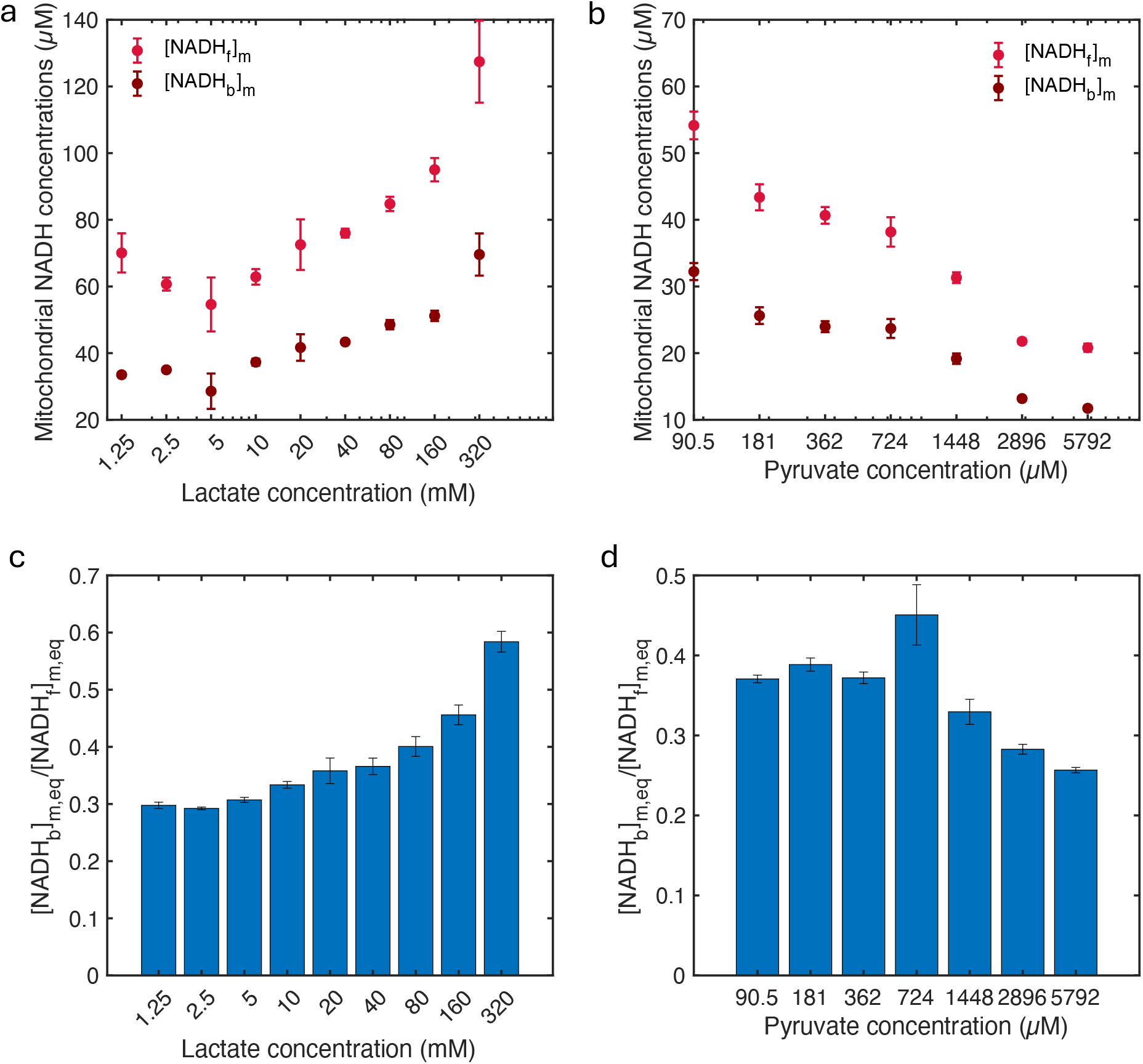
Mitochondrial NADH concentrations and equilibrium [NADH_b_]_m_/[NADH_f_]_m_ ratio during external lactate and pyruvate titration. a) Mitochondrial bound NADH ([NADH_b_]_m_) and free NADH ([NADH_f_]_m_) concentrations for titration of external lactate concentrations at fixed external pyruvate concentration of 181µM. b) Mitochondrial bound NADH ([NADH_b_]_m_) and free NADH ([NADH_f_]_m_) concentrations for titration of external pyruvate concentrations at fixed external lactate concentration of 10mM. c) Equilibrium [NADH_b_]_m,eq_/[NADH_f_]_m,eq_ obtained as the lowest [NADH_b_]_m_/[NADH_f_]_m_ ratio during oxygen drop at corresponding lactate concentrations. d) Equilibrium [NADH_b_]_m,eq_/[NADH_f_]_m,eq_ obtained as the lowest [NADH_b_]_m_/[NADH_f_]_m_ ratio during oxygen drop at corresponding pyruvate concentrations.

### Inferring oxygen consumption rate and LDH fluxes for pyruvate titration without lactate in the media

**Figure 7S.**
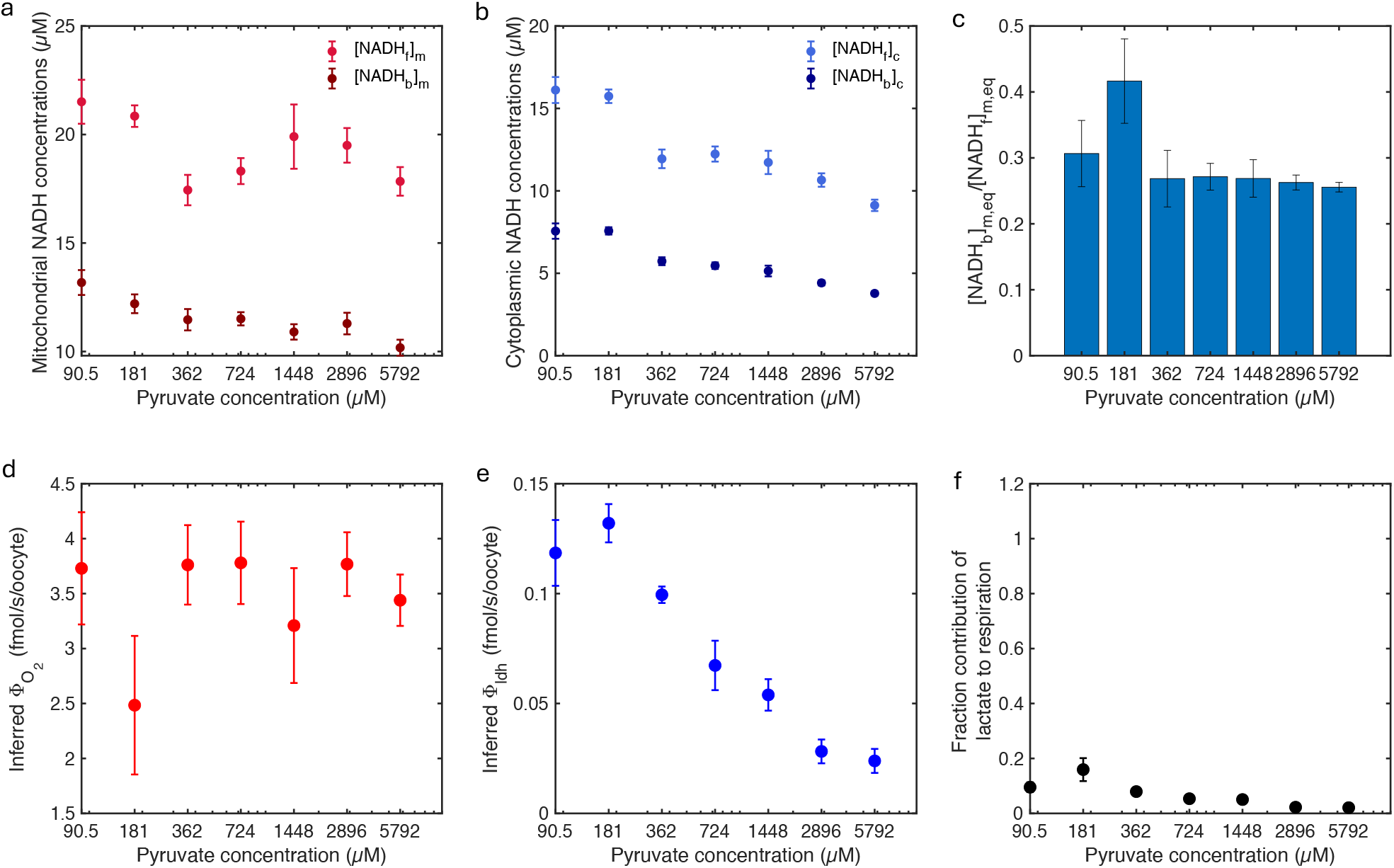
Inferring oxygen consumption rate and LDH fluxes for pyruvate titration without lactate in the media. **a)** Mitochondrial bound NADH ([NADH_b_]_m_) and free NADH ([NADH_f_]_m_) concentrations for titration of external pyruvate concentrations without external lactate. b) Cytoplasmic bound NADH ([NADH_b_]_c_) and free NADH ([NADH_f_]_c_) concentrations for titration of external pyruvate concentrations without external lactate. c) Equilibrium [NADH_b_]_m,eq_/[NADH_f_]_m,eq_ obtained as the lowest [NADH_b_]_m_/[NADH_f_]_m_ ratio during oxygen drop at corresponding pyruvate concentrations and no lactate. d) Inferred oxygen consumption rate per oocyte as a function of external pyruvate concentration without external lactate. e) Inferred LDH flux per oocyte as a function of external pyruvate concentration without external lactate. f) Fraction contribution of lactate to respiration as a function of external pyruvate concentration without external lactate.

